# *Drosophila* Snazarus regulates a lipid droplet population at plasma membrane-droplet contacts in adipocytes

**DOI:** 10.1101/620278

**Authors:** Rupali Ugrankar, Jade Bowerman, Hanaa Hariri, Mintu Chandra, Kevin Chen, Marie-France Bossanyi, Sanchari Datta, Sean Rogers, Kaitlyn M. Eckert, Gonçalo Vale, Alexia Victoria, Joseph Fresquez, Jeffrey G. McDonald, Steve Jean, Brett M. Collins, W. Mike Henne

**Author notes:** Correspondence: W. Mike Henne.

## Abstract

Adipocytes store nutrients as lipid droplets (LDs), but how they organize their LD stores to balance lipid uptake, storage, and mobilization remains poorly understood. Here, using *Drosophila* fat body (FB) adipocytes we characterize spatially distinct LD populations that are maintained by different lipid pools. We identify peripheral LDs (pLDs) that make close contact with the plasma membrane (PM) and are maintained by lipophorin-dependent lipid trafficking. pLDs are distinct from larger cytoplasmic medial LDs (mLDs) which are maintained by FASN1-dependent *de novo* lipogenesis. We find that sorting nexin CG1514/Snazarus (Snz) associates with pLDs and regulates LD homeostasis at ER-PM contact sites. Loss of *SNZ* perturbs pLD organization whereas Snz over-expression drives LD expansion, triacylglyceride production, starvation resistance, and lifespan extension through a DESAT1-dependent pathway. We propose that *Drosophila* adipocytes maintain spatially distinct LD populations and identify Snz as a novel regulator of LD organization and inter-organelle crosstalk.

## Introduction

Life presents energetic and metabolic challenges, and metazoans have developed specialized nutrient-storing organs to maintain energy homeostasis and buffer against the ever-changing availability of dietary nutrients. *Drosophila melanogaster* is a key model organism to study energy homeostasis as many aspects of mammalian metabolism are conserved in the fly (Baker and Thummel, 2007) (Lehmann, 2018) (Musselman et al., 2013). The major energy-storage organ of insects is the fat body (FB), a central metabolic tissue that exhibits physiological functions analogous to the mammalian adipose tissue and liver including nutrient storage, endocrine secretion, and immune response (Arrese and Soulages, 2010). Consequently, the FB makes intimate contact with both the gut where dietary nutrients are absorbed, and circulating hemolymph that transports lipids between organs. *Drosophila* larvae feed continuously to promote an increase in animal mass, and absorb dietary nutrients into the FB to store these as glycogen or triacylglyceride (TAG) that is incorporated into cytoplasmic lipid droplets (LDs). TAG storage ultimately requires LD biogenesis on the surface of the endoplasmic reticulum (ER), the primary site of TAG synthesis (Wilfling et al., 2014). During development or when nutrients are scarce, FB cells adapt their metabolism to mobilize LDs via cytoplasmic lipases. These mobilized lipids are delivered to other organs in the hemolymph via protein shuttles called lipophorin (Lpp) particles that are analogous to mammalian VLDL particles, but how LD mobilization is related to Lpp particle lipid loading remains poorly understood (Arrese and Wells, 1997).

The mechanisms that govern lipid flux across the FB cell plasma membrane (PM) also remain poorly characterized, but are essential for lipid export as well as lipid uptake and storage in LDs. In insects, the internalization of hemolymph lipids into both the FB and imaginal discs is unaffected when endocytosis is blocked, suggesting a non-vesicular uptake mechanism (Parra-Peralbo and Culi, 2011) (Dantuma et al., 1997) (Rodriguez-Vazquez et al., 2015). In line with this, Lpp proteins are not degraded via endolysosomal trafficking within the FB, consistent with a model where Lpp particles can donate and receive lipids directly at the FB cell surface (Dantuma et al., 1997) (Rodriguez-Vazquez et al., 2015). Furthermore, Lpp particles primarily transport diacylglyceride (DAG), suggesting Lpp-derived lipids are processed during their uptake and delivery to the ER by ER-resident acyl CoA:diacylglycerol acyltransferase (DGAT) enzymes, which convert DAG to TAG. In addition to storing extracellular Lpp-derived lipids, FB cells also generate their own lipids via fatty acid *de novo* lipogenesis. FB cells deficient in fatty acid synthesis (FAS) enzymes exhibit severe lipodystrophy, indicating FB cells somehow balance the storage of Lpp-derived and *de novo* synthesized lipids to maintain fat homeostasis (Wicker-Thomas et al., 2015) (Heier and Kuhnlein, 2018).

Due to their specialized function in lipid uptake and storage, many fat-storing cells exhibit a unique surface architecture: their PM is densely pitted with invaginations that increase the surface area exposed to the extracellular space (Diaconeasa et al., 2013) (Pilch and Liu, 2011). In mammals, up to half the surface of white adipocytes is decorated with caveolae, invaginations that organize surface receptors as well as promote lipid and nutrient absorption (Pilch et al., 2011). Surprisingly, *Drosophila* do not encode caveolin genes that are required to form caveolae (Gupta et al., 2009). Nevertheless, *Drosophila* FB adipocytes exhibit their own intricate networks of surface invaginations that are stabilized by the cortical actin network (Diaconeasa et al., 2013). Perturbing this cortical actin network disrupts FB lipid homeostasis, suggesting a functional connection between FB surface architecture and lipid storage (Mazock et al., 2010) (Diaconeasa et al., 2013).

Although LDs serve as organelle-scale lipid reservoirs, how cells organize their LD stores to balance storage with efficient mobilization is largely unresolved (Thiam and Beller, 2017) (Zhang et al., 2016). An intuitive mechanism to organize LDs is to attach them to other organelles, as this allows them to exchange lipids with these organelles as well as potentially compartmentalize them in distinct regions of the cell interior (Olzmann and Carvalho, 2019) (Henne et al., 2018). Recent work using *Saccharomyces cerevisiae* reveal that even simple yeast contain functionally distinct LD sub-populations that are spatially compartmentalized. This compartmentalization is achieved by LD-organizing proteins that bind to LDs and cluster them adjacent to the yeast vacuole/lysosome (Eisenberg-Bord et al., 2017) (Teixeira et al., 2017) (Hariri et al., 2018). One such organizing protein is Mdm1, an ER-anchored protein that binds to LDs and attaches them to the vacuole/lysosome surface (Henne et al., 2015) (Hariri et al., 2018). Mdm1 is highly conserved in *Drosophila* as CG1514/Snazarus (Snz), originally characterized as a longevity-associated gene of unknown function that is highly expressed in the *Drosophila* FB (Suh et al., 2008). Both yeast and human Snz homologs bind to LDs and regulate LD homeostasis, but the function of Snz in *Drosophila* remains unclear (Hariri et al., 2019) (Datta et al., 2019).

Here, we investigate how FB cells functionally and spatially organize their LD stores. We find that FB cells contain functionally distinct LD populations that are spatially segregated into regions of the cell interior. These LD populations require distinct lipid pools for their maintenance, with LDs in the cell periphery (peripheral LDs, pLDs) requiring Lpp-dependent trafficking, whereas LDs further in the cell interior (medial LDs, mLDs) are maintained by FASN1-dependent *de novo* lipogenesis within the FB. We also characterize Snz as a novel regulator of pLD homeostasis that localizes to ER-PM contacts and promotes LD growth and TAG storage.

## Results

### *Drosophila* FB cells contain small peripheral LDs (pLDs) wrapped by plasma membrane invaginations

To investigate how *Drosophila* FB adipocytes functionally organize their LD stores, FB tissue was extracted from feeding L3 larvae and imaged using high-resolution fluorescence confocal microscopy. Extracted FB tissue was fixed and stained with the fluorescent LD dye monodansylpentane (MDH), then imaged by longitudinal serial *z*-sectioning at 1 µm step-sizes. Consistent with previous studies, serial imaging revealed that FB tissue is composed of a continuous monolayer of polygonal cells ~40 µm in height. (Arrese and Soulages, 2010). Confocal sections also revealed that individual FB cells contain a range of differently sized LDs in distinct regions of the cytoplasm. In general, LDs in the cell periphery immediately below the tissue surface were small (~5μm^2^ in cross-sectional area) compared to LDs adjacent to the nucleus at the tissue mid-plane (~28μm^2^ in area) (Figure 1A-D, **SFigure 1A**). To observe LDs at higher resolution, we also conducted thin-section transmission electron microscopy (TEM) of intact FB tissue. Similar to light microscopy, TEM imaging confirmed the presence of large medial LDs (mLDs) in the tissue peri-nuclear region, and smaller peripheral LDs (pLDs) immediately below the cell surface (Figure 1E, **red arrow**).

**Figure 1:**
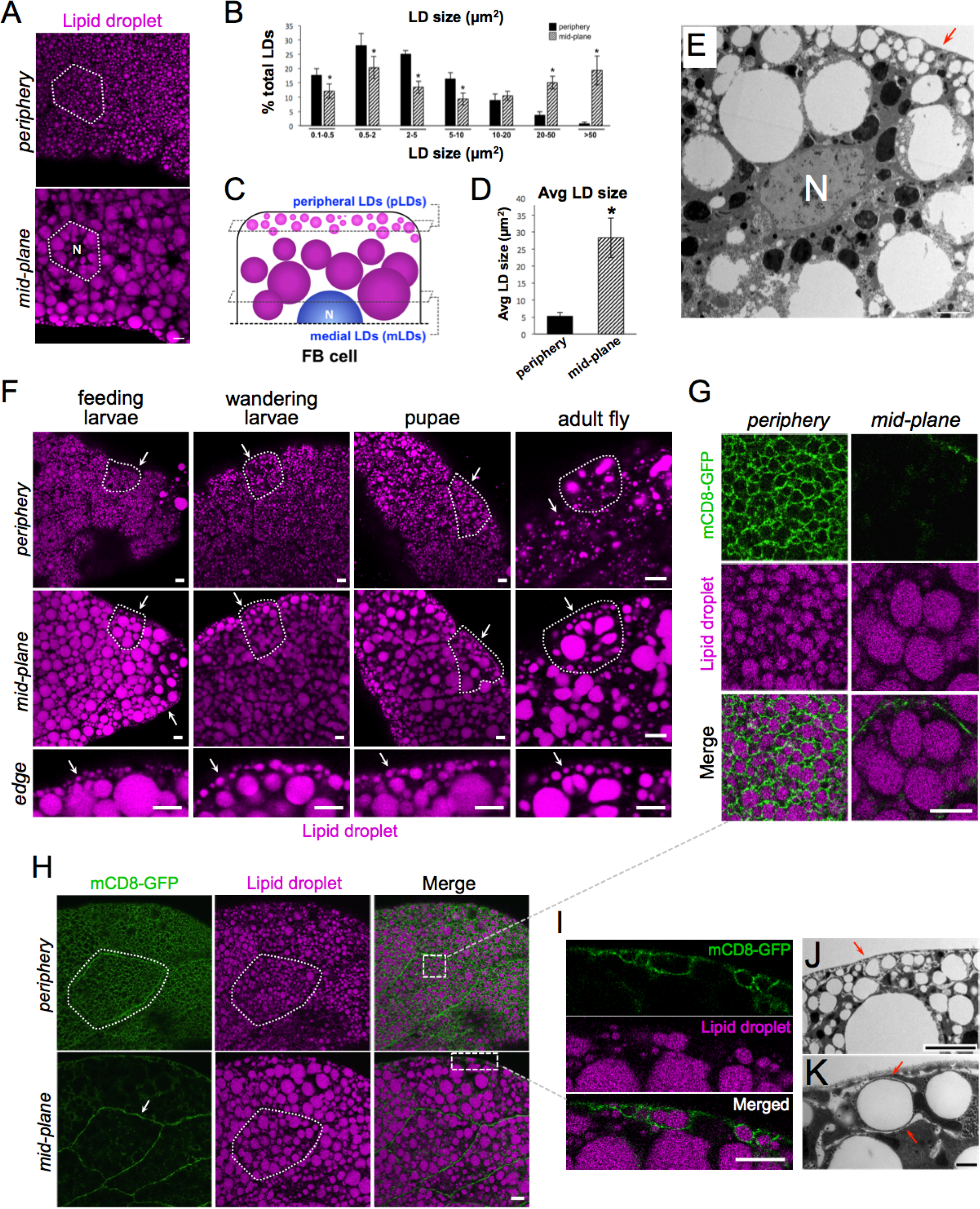
*Drosophila* FB cells contain peripheral LDs that associate with the PM. **A**) Confocal micrographs of intact larval FB tissue stained with LD marker MDH (magenta) and imaged at the tissue surface (*periphery*) and cell interior at the peri-nuclear region (*mid-plane*). Nucleus is denoted with “N”. A single FB cell is outlined in white. Scale bar 10µm. **B**) Quantification of LD sizes (LD cross-sectional areas in µm^2^) in periphery and mid-plane. **C**) Cartoon summarizing peripheral LDs (pLDs) and medial LDs (mLDs). **D**) Average LD size in periphery and mid-plane. **E**) TEM micrograph of FB cell mid-plane showing peripheral LD (pLDs, red arrow) and larger mLDs. Nucleus is denoted (N). Scale bar 10µm. **F**) FB from different stages of *Drosophila* development and stained with MDH. Cell periphery and mid-plane is shown for each tissue and arrows denote the pLD layer. A single cell is outlined in each tissue. The tissue edge denoted the tissue surface. **G**) Zoom-in of the FB cell periphery from Panel H expressing plasma membrane (PM) marker mCD8-GFP (*Dcg-Gal4>UAS-mCD8-GFP*) and stained with MDH. **H**) FB expressing mCD8-GFP and MDH stain. Single cell is outlined and denoted. Boxed insets denote regions shown in Panels G and I. **I**) Zoom-in of cell edge with mCD8-GFP-positive PM wrapping around pLDs. **J**) TEM micrograph of FB periphery showing tubular membranes in cell periphery and their close association with pLDs (red arrows). **K**) TEM micrograph showing tubular wrapping around pLD. Scale bar 1µm. Scale bar is 10µm unless otherwise denoted.

To determine whether these LD populations existed throughout the different stages of *Drosophila* development, we examined FB tissue from feeding L3 larvae, late wandering L3 larvae preparing to pupate, and early stage pupae. FBs from all stages contained small pLDs that formed a distinct pLD layer immediately below the tissue surface (Figure 1F, **arrows**). We also examined adult FB cells from 1-week-old adult files. Larval FB cells are actively turned over within the first few days of fly emergence, and adults generate new FB cells *de novo* (Arrese and Soulages, 2010). Similar to larval fat cells, adult adipocytes also contained small pLDs near the tissue surface, although adult LDs were less densely packed than in larvae (Figure 1F).

We next examined the association of pLDs to the cell plasma membrane (PM). We expressed the PM marker mCD8-GFP in FB cells using the FB tissue-specific driver *Dcg-Gal4* (*Dcg-Gal4>UAS-mCD8-GFP*) (Suh et al., 2008). Strikingly, mCD8-GFP labeling revealed that the cell surface was densely ruffled, giving the PM a “foamy” appearance as previously reported (Diaconeasa et al., 2013) (Figure 1G,H, **SMovie 1**). Surprisingly, these PM ruffles were closely associated with pLDs and appeared to wrap around them, suggesting the PM and pLDs were in close physical contact (Figure 1G,H,I). We confirmed that this PM wrapping was not unique to the mCD8-GFP marker by also labeling the PM with the PtdIns(4,5)P_2_-binding PLCδ-2×PH domain (*Dcg-Gal4>mCherry-PLCδ-2×PH*) (**SFigure 1B**). We also detected PM associations with pLDs in adult FB cells (**SFigure 1C**). To evaluate the PM and pLDs at higher resolution, we conducted high-magnification TEM. Indeed, TEM revealed that the cell periphery contained numerous tubular structures that closely wrapped around pLDs, often surrounding pLDs and creating ~10nm of space between the tubular membrane and the pLD surface (Figure 1J,K, **SFigure 1D,E**). Collectively, we conclude that *Drosophila* FB cells contain large mLDs and small pLDs, the latter making close contact with the cell surface.

### pLDs change in response to alterations in organismal nutrient status and lipophorin trafficking

Since pLDs establish close contact with the PM, we hypothesized that pLDs may serve as lipid reservoirs that store lipids either derived from or destined for the extracellular hemolymph. To investigate this possibility, *w*^1118^ control larvae were fasted for 24 hours to induce LD mobilization. Fasting induced significant changes to pLD morphology. There were reduced numbers of pLDs leading to a significant decrease in pLD density from ~105 pLDs down to ~67 pLDs per 900 µm^2^ (Figure 2A,C). There was also a corresponding increase in remaining pLD size from ~3 µm^2^ to ~7 µm^2^ (Figure 2A,B). In contrast, fasting induced no significant changes to the mLD population; average mLD size and density were similar in fed and fasting states, indicating fasting primarily affected the pLD population (**SFigure2 A,B**).

**Figure 2:**
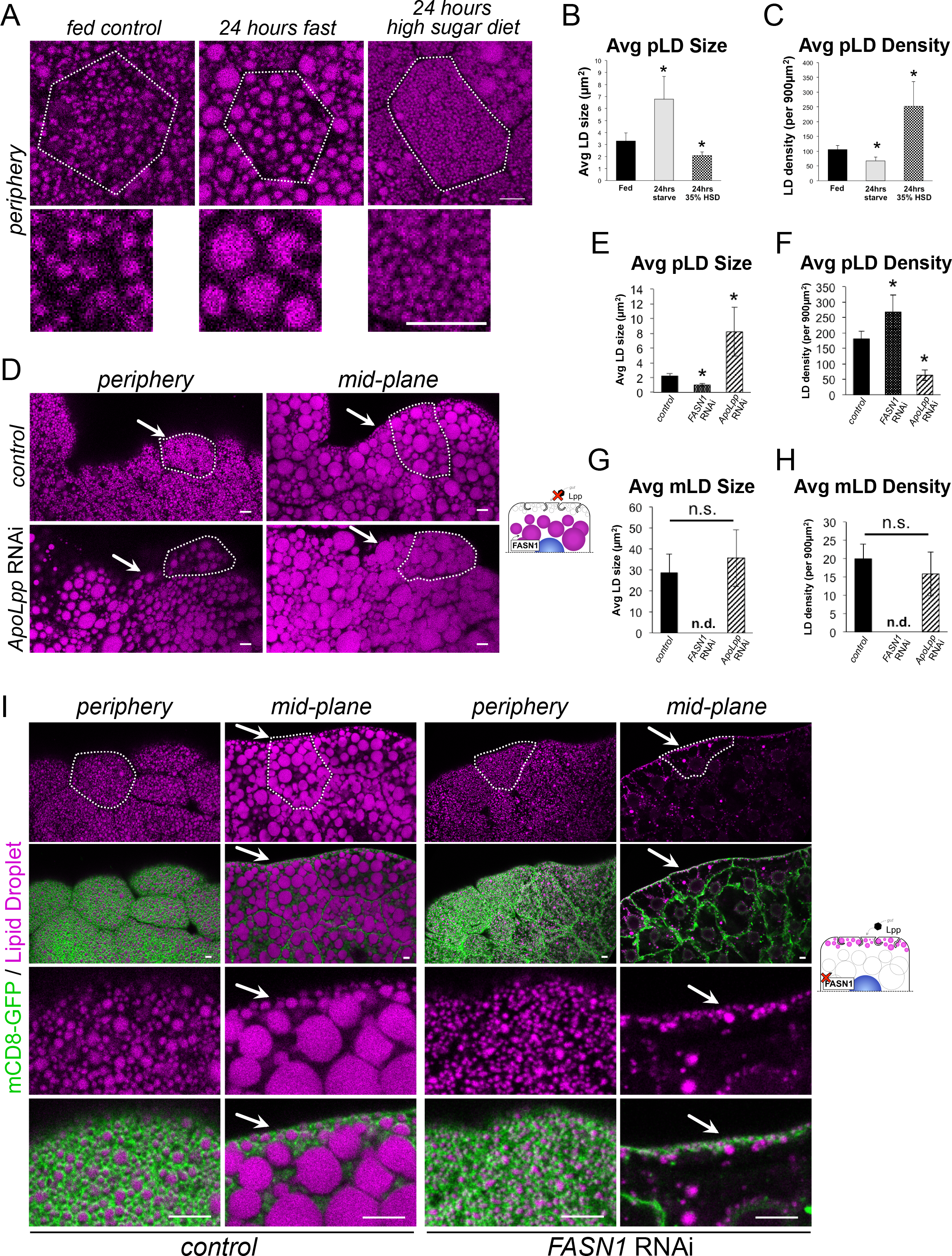
FB pLDs and mLDs are maintained by distinct lipid sources. **A**) FB cell peripheries stained for LDs (MDH, magenta) in either fed larvae, 24 hour fasted larvae, or larvae fed a high sugar diet for 24 hours. White outline indicates a single cell boundary. Scale bars 10µm. **B**) Quantification of average LD size of samples in Panel A. * indicates p<0.05. **C**) Average LD density for Panel A. * indicates p<0.05. **D**) FB tissue stained for LDs (MDH, magenta) in either control (*Dcg-Gal4*) or (*Dcg-Gal4>UAS-ApoLpp*^*RNAi*^). Arrows indicate areas the pLDs should reside. Scale bar is 10µm. **E**) Average pLD size for control (*Dcg-Gal4*), *FASN1^RNAi^ (Dcg-Gal4>UAS-FASN1^RNAi^)*, and *ApoLpp*^RNAi^ samples. * indicates p<0.05. **F**) Average pLD density for samples as in E. **G**) Average mLD size for control (*Dcg-Gal4*), *FASN1*^*RNAi*^, and *ApoLpp*^RNAi^ samples. * indicates p<0.05. **H**) Average mLD density for samples as in G. **I**) FB stained for LDs (MDH, magenta) and expressing mCD8-GFP PM marker (*Dcg-Gal4>UAS-mCD8-GFP*) (green). Individual cells are outlined in white. Control (*Dcg-Gal4*) samples are on the left and *FASN1*^*RNAi*^ FB samples are on the right. Arrows indicate the pLD population. Scale bar is 10µm.

Next, *w*^1118^ larvae were reared for 24 hours on a high-sugar diet (HSD, 35% sugars), which stimulates lipogenesis in several organs including the gut and promotes lipid trafficking in the hemolymph and delivery to the FB for storage (Palm et al., 2012) (Musselman et al., 2011). HSD also significantly altered pLDs; there was a striking boost in the number of very small pLDs (<2 µm^2^) near the cell surface, equating to a decrease in average pLD size (Figure 2A,B). This accumulation of small pLDs led to a more than 2-fold increase in pLD density from ~105 pLDs to ~252 pLDs per 900 µm^2^ (Figure 2A, C). There were also moderate changes to the mLD population. mLD size became more variable as small LDs now appeared at the tissue edges in the cell mid-plane (**SFigure 2A**). There was also a moderate but significant decrease in mLD density from ~20 mLDs to ~14 mLDs per 900 µm^2^ (**SFigure 2B**).

Collectively we conclude that changes in larval dietary status differentially affect the pLD and mLD populations. Fasting primarily affects the pLD population, leading to decreased pLD abundance and an increase in the sizes of the remaining pLDs. Notably, mLD sizes and densities are not altered during fasting. In contrast, HSD feeding results in greater numbers of densely packed pLDs near the cell surface, and mLDs become more heterogeneous with small LDs now infiltrating into the mLD layer.

### pLDs and mLDs are differentially maintained by lipophorin trafficking and FB *de novo* lipogenesis

We hypothesized that pLDs may contain lipids derived from lipophorin (Lpp) particles, which transport dietary lipids from gut enterocytes through the hemolymph to organs including the FB for storage (Arrese et al., 2001). To investigate this, we RNAi-depleted Lpp particles by knocking down the major Lpp structural protein ApoLpp (*Dcg-Gal4>UAS-ApoLpp^RNAi^*), which is expressed almost exclusively in the FB from where it is secreted into the hemolymph (Palm et al., 2012). *ApoLpp^RNAi^* larvae were viable but displayed developmental delay and were unable to develop into adult flies, as previously reported (Palm et al., 2012). Intriguingly, *ApoLpp*^*RNAi*^ drastically affected the pLD population. pLDs were more sparse, and FB cells manifested a more than 50% reduction in pLD density (Figure 2D, F). Furthermore, the remaining LDs near the cell surface exhibited a more than 3-fold increase in size, and appeared similar to mLDs (Figure 2D-F, **SFigure 2C**). As a consequence, the normally distinct layer of pLDs lining the cell periphery was absent in *ApoLpp*^*RNAi*^ FBs (Figure 2D, **SFigure 2C**). In line with this, mCD8-GFP-expressing *ApoLpp*^*RNAi*^ cells exhibited reduced PM association with LDs (**SFigure 2F**). In contrast, the mLD population was generally unaffected in *ApoLpp*^*RNAi*^ cells, and no significant change was manifested in average mLD size nor density (**SFigure 2D,E**). This is consistent with previous work indicating that the bulk of TAG in the FB is derived from intracellular *de novo* lipogenesis (Palm et al., 2012). Collectively, this suggests that the FB pLD population requires some aspect of Lpp trafficking to be properly maintained near the cell surface.

Next we determined whether the pLD and mLD populations were affected by perturbation of *de novo* lipogenesis within the FB. *Drosophila* encode three fatty acid synthase (FASN) proteins that are required for *de novo* lipogenesis, with FASN1 responsible for most FA synthesis within the FB (Parvy et al., 2012) (Garrido et al., 2015). We knocked-down *FASN1* specifically in the FB tissue by RNAi (*Dcg-Gal4>UAS-FASN1^RNAi^*), and examined larval FBs. Notably, *FASN1^RNAi^*-treated larvae were viable but exhibited drastic lipodystrophy and the extracted FBs sank to the bottom of PBS-containing reservoirs as they were gathered for experiments. In line with this, *FASN1*^*RNAi*^ FBs were largely devoid of intracellular mLDs, with only a few small LDs remaining in the cell interior (Figure 2G, H, I). However, strikingly the pLD population remained intact immediately beneath the cell surface, and the cell periphery exhibited a dense array of pLDs adjacent to the PM that were similar to pLDs in control FBs (Figure 2I). In fact, pLD density actually increased slightly in *FASN1*^*RNAi*^ tissues, although average pLD size decreased slightly (Figure 2E,F). Consistent with this, *FASN1*^*RNAi*^ larvae exhibited a more than 80% reduction in whole-organism TAG. In contrast, *ApoLpp*^*RNAi*^ larvae had only minor changes in global TAG levels, although there was a significant reduction in organismal DAG consistent with a loss of Lpp-associated DAG pools (**SFigure 2G-J**). Collectively, this suggests that mLDs require FASN1-dependent *de novo* lipogenesis in the FB to be properly maintained. In contrast, the pLD population is independent of FASN1-mediated lipogenesis within the FB, but rather relies on some aspect of Lpp trafficking for their maintenance.

### pLDs are maintained by a cortical actin network and perilipin LSD2

We next investigated what structural factors maintain the pLD layer adjacent to the cell surface. First, we disrupted the FB cell PM invaginations by tissue-specific RNAi-depletion of the cortical actin scaffolding protein β-spectrin (*Dcg-Gal4>UAS-β-spectrin*^*RNAi*^), which was previously shown to be required for maintaining the “foamy” appearance of the cell surface (Figure 3A, B) (Diaconeasa et al., 2013) As expected, loss of β-spectrin caused a near complete loss of PM invaginations. Interestingly, this also perturbed pLD morphology and density in the cell periphery (Figure 3A, B). pLDs now appeared disorganized and became more varied in size, reflecting an increase in average pLD size but a decrease in pLD density (Figure 3C,D). In contrast, mLD size and density were unaffected by knock down of β-spectrin, indicating this affected primarily the pLD sub-population (**SFigure 3A,B**). This suggests that a cortical actin network is required to maintain the FB cell PM architecture as well as maintain regular pLD morphology, however mLDs do not require this.

**Figure 3:**
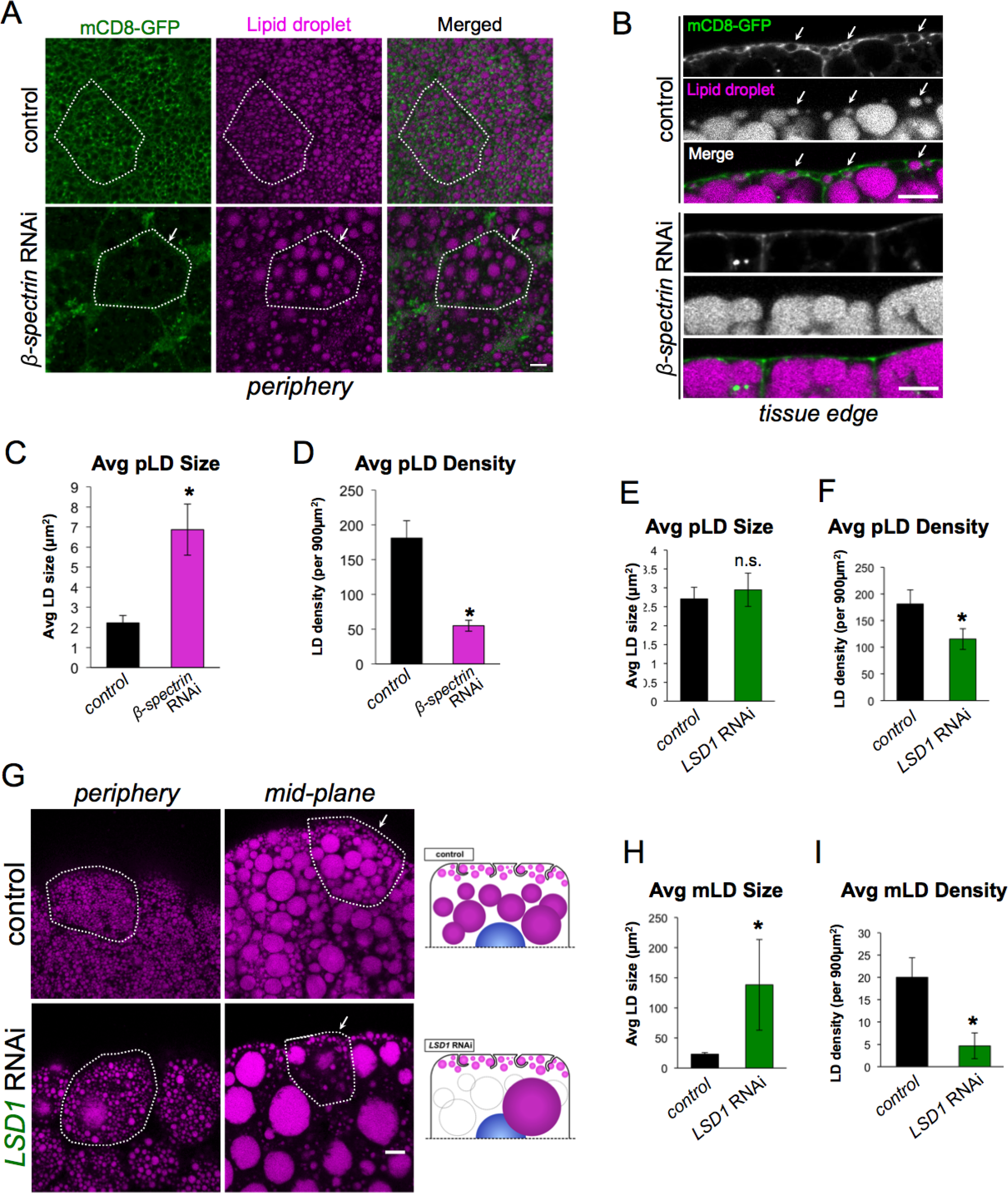
pLD morphology is maintained by β-spectrin and LSD2 but is independent of perilipin LSD1. **A**) FB expressing mCD8-GFP PM marker (green) (*Dcg-Gal4>UAS-mCD8-GFP*) and stained for LDs (MDH, magenta). Tissues are either control (*Dcg-Gal4* alone) or β*-spectrin*^RNAi^ (*Dcg-Gal4>UAS-β-spectrin*^*RNAi*^) in the FB. Individual cells are outlined in white. Arrows indicate a cell with altered pLD morphology. Scale bar is 10µm. **B**) Side profiles of control and *β-spectrin*^RNAi^ tissue as in Panel A. White arrows indicate PM-wrapping of pLDs in control, which are missing in *B-spectrin*^RNAi^ tissue. Scale bar is 10µm. **C**) Average pLD size in control and *β-spectrin*^RNAi^ tissue. * indicates p<0.05. **D**) Average pLD density in control and *β-spectrin*^RNAi^ tissue. * indicates p<0.05. **E**) Average pLD size in control (*Dcg-Gal4*) and *LSD1*^RNAi^ (*Dcg-Gal4>UAS-LSD1*^*RNAi*^) tissue. **F**) Average pLD density in control (*Dcg-Gal4*) and *LSD1*^RNAi^ tissue. * indicates p<0.05. **G**) FB stained for LDs (MDH, magenta) of either control (*Dcg-Gal4* alone) or *LSD1*^RNAi^. Single cells are outlined in white. Arrows indicate the pLD layer. Scale bar is 10µm. **H**) Average mLD size in control and *LSD1*^RNAi^ tissue. * indicates p<0.05. **I**) Average mLD density in control (*Dcg-Gal4*) and *LSD1*^RNAi^ tissue. * indicates p<0.05.

Next, we evaluated whether pLDs and mLDs were differentially dependent on the perilipin proteins that coat LD surfaces. *Drosophila* encode two perilipin proteins: LSD1 and LSD2. Previous studies indicated that these are differentially targeted to LDs, with LSD1 localizing to the surfaces of all LDs, whereas LSD2 is enriched primarily on small LDs in the FB (Bi et al., 2012). Intriguingly, FB-specific RNAi-depletion of LSD1 (*Dcg-Gal4>LSD1*^*RNAi*^) led to a drastic increase in the size of individual mLDs, and a corresponding reduction in the number of mLDs in the peri-nuclear region (Figure 3G, H, I). In contrast, the pLD population was largely unaffected by loss of *LSD1*, and a pLD population was still very clearly distinguishable below the cell surface in *LSD1*^*RNAi*^ FBs (Figure 3G). In line with this, average pLD size was not affected in *LSD1*^*RNAi*^ FB cells, although there was a slight decrease in pLD surface density (Figure 3E,F).

Next we RNAi-depleted *LSD2* in the FB (*Dcg-Gal4>UAS-LSD2*^*RNAi*^). In contrast to LSD1, *LSD2*^*RNAi*^ failed to alter the mLD population (**SFigure 3C,F,G**). However, the pLD population was now affected and exhibited an increase in average pLD size and decrease in pLD density (**SFigure 3C-E**). Collectively, this indicates that LSD1 and LSD2 are differentially required to maintain the morphologies of the mLD and pLD populations. LSD1 appears necessary to maintain the regular morphology of the mLD population, but is dispensable for the maintenance of the pLD population. In contrast, LSD2 is required to maintain the regularity of the pLD population. Indeed, *LSD2*^*RNAi*^ FB cells contain morphologically heterogeneous pLDs that appear similar to pLDs in *β-spectrin*^*RNAi*^ tissue.

### Snazarus (Snz) is an ER-associated protein that enriches at ER-PM contacts

As FB cells contained distinct LD populations, we next investigated whether proteins known to associate with distinct LD populations in other organisms exhibited any association with LDs in the FB. Recent studies in budding yeast have highlighted numerous LD binding proteins that control the spatial distribution of specific LD populations within the cytoplasm (Henne et al., 2018) (Olzmann and Carvalho, 2019). Many of these “LD organizing” proteins have clear homologs in *Drosophila*, and one such homolog is CG1514/Snazarus (Snz). Snz is a poorly characterized protein and clear homolog of yeast Mdm1, an inter-organelle tether that binds to LDs and clusters them adjacent to the yeast vacuole (Hariri et al., 2018) (Hariri et al., 2019). Mdm1 is an ER-resident protein that binds to LDs as they bud from the ER surface. Because Mdm1 also contains a C-terminal PtdIns(3)P-binding Phox homolog (PX) domain that attaches to the yeast vacuole, it can promote the accumulation of LDs adjacent to the vacuole at the ER-vacuole interface (Hariri et al., 2019). Snz features the same domain architecture as Mdm1, suggesting a conservation of function (Suh et al., 2008) (Henne et al., 2015). Furthermore, previous work and mRNA datasets indicate that Snz is highly expressed in *Drosophila* FB tissue (Suh et al., 2008). We also confirmed by qRT-PCR that Snz is expressed in both larvae and adult flies (**SFigure 4A**).

**Figure 4:**
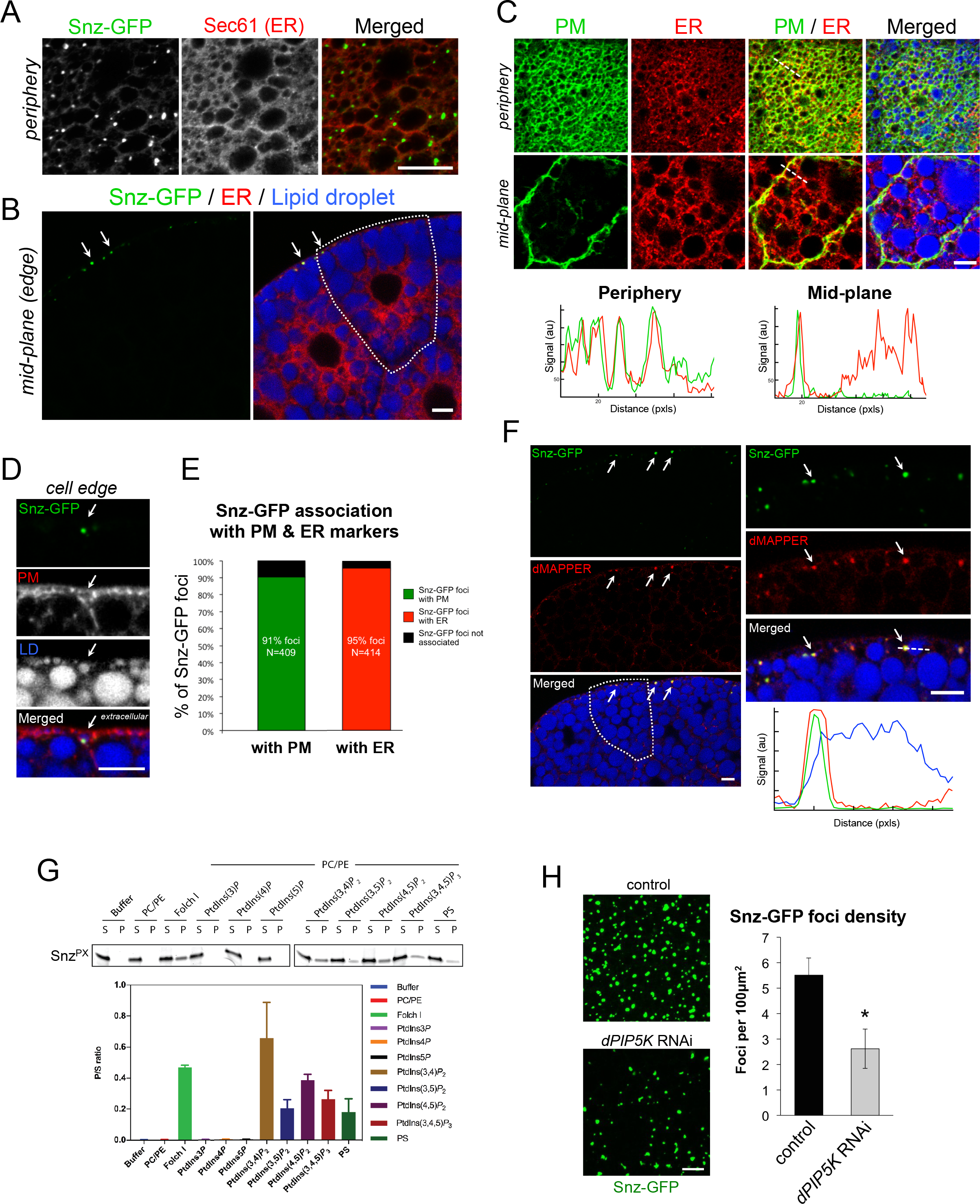
Snz is an ER-associated protein that enriches at ER-PM contact sites. **A**) Immuno-fluorescence stained FB tissue periphery expressing Snz-GFP (*Dcg-Gal4>UAS-Snz-GFP*) (green) that forms foci that co-localize along the ER network (anti: Sec61, red). Scale bar 10µm. **B**) FB tissue expressing Snz-GFP (green), ER marker Sec61-TdTomato (red) (*Dcg-Gal4>UAS-Sec61-TdTomato*), and stained for LDs (MDH, blue). A single cell is outlined in white. Arrows denote Snz-GFP foci at the cell periphery. Scale bar 10µm. **C**) FB tissue expressing PM marker mCD8-GFP (green) (*Dcg-Gal4>UAS-mCD8-GFP*) and ER marker Sec61-TdTomato (red), and stained for LDs (MDH, blue). Line-scans noted with white line. Scale bar 10µm. **D**) Side profile of FB tissue periphery expressing Snz-GFP (green) (*Dcg-Gal4>UAS-Snz-GFP*) and PM marker mCD8-mCherry (red) (*Dcg-Gal4>UAS-mCD8-mCherry*), and LD stain (MDH, blue). Arrow indicates a Snz-GFP punctae associated with a PM-wrapped pLD. Scale bar 10µm. **E**) Quantification of Snz-GFP foci co-localizing with either PM marker (mCD8-mCherry) or ER marker (Sec61-TdTomato) from independent FB imaging experiments. **F**) FB tissue periphery expressing Snz-GFP (green) (*Dcg-Gal4>UAS-Snz-GFP*), dMAPPER (red) (*Dcg-Gal4>UAS-dMAPPER-mCherry*), and stained for LDs (MDH, blue). Arrows indicate examples of Snz-GFP and dMAPPER co-localization. Linescan denoted with white line. Scale bar 10µm. **G**) Liposome co-sedimentation assay using recombinant *E. coli* purified Snz PX domain and liposomes containing different phospholipid compositions. Ratio of liposome bound (pellet, P) to unbound (supernatant, S) is graphed. **H**) FB cell peripheries expressing Snz-GFP in either control (*Dcg-Gal4* alone) or dPIP5K^RNAi^-treated (*Dcg-Gal4>dPIP5K*^*RNAi*^) FB tissue. Quantification of Snz-GFP foci density is graphed. * indicates p<0.05. Scale bar 10µm unless otherwise indicated.

To determine the sub-cellular localization of Snz, we imaged FB tissue expressing a C-terminally GFP-tagged Snz (Snz-GFP) (*Dcg-gal4>UAS-Snz-GFP*). Snz-GFP was detected as distinct intra-cellular foci (Figure 4A). Since Snz contains a predicted N-terminal transmembrane region, we interrogated whether Snz-GFP was ER associated like its yeast and mammalian orthologs (Henne et al., 2015) (Bryant et al., 2018) (Datta et al., 2019). Co-immunofluorescence revealed that Snz-GFP co-localized with ER protein Sec61 along the ER tubular network, and dimmer Snz-GFP signal could also be observed along ER tubules, suggesting an ER association (Figure 4A). We also co-stained the FB with ER marker Calnexin 99a that confirmed the Snz-GFP ER association (**SFigure 4B**).

Remarkably, z-section confocal imaging revealed that Snz-GFP foci were highly polarized to the FB cell periphery, and were generally absent from the cell interior (Figure 4B). Consistent with this, stacked z-sections through the FB tissue revealed that Snz-GFP foci were polarized to the extreme apical and basal tissue surfaces, as well as along cell edges exposed to the hemolymph (**SFigure 4C-F**). Furthermore, the Snz-GFP foci were positioned in the same tissue layer as the pLDs, with ~65% of foci within 5µm of the surface layer of the FB (**SFigure 4E, F**).

To more directly evaluate the spatial relationship between the PM and Snz-GFP foci, we imaged FB tissue co-expressing Snz-GFP and the PM marker mCD8-mCherry (*Dcg-Gal4>UAS-Snz-GFP; Dcg-Gal4>UAS-mCD8-mCherry*). Remarkably, Snz-GFP foci co-localized with mCD8-mCherry, and often localized specifically with PM regions that were wrapping around pLDs (Figure 4D, **SFigure 4G**). Quantification of 409 Snz-GFP foci showed that 91% co-localized with the mCD8-mCherry PM signal (Figure 4E). Since Snz-GFP foci were also ER-associated we also imaged tissue co-expressing Snz-GFP and Sec61-TdTomato (*Dcg-Gal4>UAS-Snz-GFP; Dcg-Gal4>UAS-Sec61-TdTomato*). Indeed, 95% of 414 Snz-GFP foci co-localized with Sec61-TdTomato-positive ER tubular structures (Figure 4E, **SFigure 4G**). Consistent with this, co-labeling both the PM (mCD8-GFP) and ER (Sec61-TdTomato) together revealed that the PM and ER network were closely overlaid in the cell periphery (Figure 4C). Line-scans confirmed ER and PM signal overlap in the periphery but not the cell mid-plane, suggesting the ER and PM are in close proximity (Figure 4C). This was further supported by TEM imaging which revealed extensive networks of ER tubules near the cell surface (**SFigure 1C,D, red arrows**).

Given its co-localization with both ER and PM markers, we hypothesized that Snz-GFP foci were enriching at sites of close juxtaposition between the ER and PM at so-called ER-PM contact sites. To test this, we generated a *Drosophila* line encoding the ER-PM contact site biomarker MAPPER (dMAPPER), which is anchored in the ER via a transmembrane domain and spans the cytoplasm to bind *in trans* to electrostatically charged lipids on the PM (Chang et al., 2013). Expression of dMAPPER-mCherry formed distinct foci in the cell periphery that co-localized with Snz-GFP foci (*Dcg-Gal4>UAS-Snz-GFP; Dcg-Gal4>UAS-dMAPPER-mCherry*) (Figure 4F). Collectively, this suggests that: 1) *Drosophila* FB cells contain regions of close ER and PM coupling which can be labeled with dMAPPER, and 2) Snz-GFP foci reside at the cell periphery and enrich at dMAPPER-positive regions of close contact between the ER and PM.

### Snz binds phospholipids via a non-canonical Phox homology (PX) domain

The co-localization of Snz-GFP foci with dMAPPER and PM markers suggested Snz may directly bind to the PM *in trans* from the ER. This would be topologically similar to how Mdm1 acts as an inter-organelle tether between the ER and yeast vacuole (Henne et al., 2015). To investigate whether Snz utilizes a similar mechanism, we first determined whether it required its C-terminal phospholipid-binding PX domain to form foci at the cell periphery. We expressed a GFP-tagged Snz construct lacking the C-terminal region including the PX domain (*Dcg-Gal4>UAS-Snz^ΔC^-GFP*). Snz^ΔC^-GFP failed to form foci in the FB cell periphery, and instead distributed dimly throughout the cell interior, indicating the C-terminal region is necessary for foci formation (**SFigure 4H**).

Next we expressed and purified the Snz PX domain from *E. coli* and incubated it with liposomes of distinct lipid compositions. Whereas Snz PX domain remained soluble in isolation and did not bind uncharged liposomes composed of phosphatidylcholine (PC) and phosphatidylethanolamine (PE), it co-pelleted with Folch I fraction liposomes that are rich in polar lipids (Figure 4G). Surprisingly, Snz PX failed to co-pellet with PtdIns(3)P liposomes nor with other liposomes containing the mono-phosphorylated phospholipids PtdIns(4)P or PtdIns(5)P, suggesting it exhibited a different phospholipid binding specificity than its yeast homolog Mdm1 (Figure 4G). In line with this, Snz PX co-pelleted with liposomes containing di-phosphorylated phosphoinositides including PtdIns(3,4)P_2_ and PtdIns(4,5)P_2,_ which are known to accumulate at the cell PM. Snz PX also interacted with PtdIns(3,5)P_2_ and the tri-phosphate lipid PtdIns(3,4,5)P_3_, suggesting it exhibits broad specificity for di- and tri-phosphorylated phospholipids. It also co-pelleted with phosphatidylserine (PS) containing liposomes, an electrostatically charged phospholipid also enriched on the PM (Figure 4G). This suggests that Snx PX exhibits broad phospholipid-binding specificity and can interact with negatively-charged phospholipids but does not bind to PtdIns(3)P like yeast Mdm1.

To better understand how the Snz PX domain interacts with phospholipids, we compared its sequence to mammalian Snz homologs SNX13 and SNX25 whose lipid binding properties have been structurally and biochemically well characterized (Mas et al., 2014) (Chandra M, 2019). Intriguingly, Snz PX showed strong primary sequence similarity to SNX25 which also binds di- and tri-phosphorylated phosphoinositide phospholipids (**SFigure 4I**) (Chandra M, 2019). To better understand this similarity, we modeled the Snz PX domain structure using the recently solved NMR structure of human SNX25 PX domain as a template (PDB ID: 5WOE) (**SFigure 4I**). Modeling revealed a positively charged surface on Snz PX similar to the non-canonical lipid-binding surface of SNX25 PX that mediates di-/tri-phosphorylated phospholipid binding (**SFigure 4I**). In line with this, sequence comparisons revealed that the canonical PtdIns(3)P binding site, which requires a strict set of side-chains (Arg-Tyr-Lys-Arg), was not conserved in either SNX25 or Snz (red residues), but positively-charged resides required to form the non-canonical binding surface were conserved between SNX25 and Snz PX (blue residues) (**SFigure 4I, bottom alignment**) (Mas et al., 2014) (Chandra and Collins, 2018) (Chandra M, 2019). Based on these similarities we mutated residues on the non-canonical charged surface of Snz PX domain, and conducted Folch I fraction liposome binding experiments. Whereas wildtype Snz PX domain bound Folch I fraction liposomes, there was reduced liposome binding when residues R725 and K726 on the non-canonical surface were mutated (R725A/K726A) (**SFigure 4J**). In addition, triple mutation of R728A/H729A/K735A on the Snz PX domain caused complete attenuation of liposome binding, suggesting this non-canonical charged surface was required for membrane interaction (**SFigure 4J,K**). We conclude that the Snz PX domain interacts with broad specificity with charged phospholipids via a non-canonical charged surface similar to the SNX25 PX domain.

Since PtdIns(4,5)_2_ is the most abundant charged phosphoinositide phospholipid at the PM, we RNAi depleted the PI(4)5-kinase dPIP5K that generates PtdIns(4,5)P_2_ in the FB (*Dcg-Gal4>UAS-PIP5K59B*^*RNAi*^), and determined whether this affected Snz-GFP foci (Idevall-Hagren and De Camilli, 2015). Knock-down of *dPIP5K* resulted in a ~50% loss of Snz-GFP foci density at the cell periphery (Figure 4H). We confirmed that *dPIP5K*-depleted FB cells still contained PM invaginations, indicating that the change in Snz-GFP foci density was not due to a loss of surface architecture (**SFigure 4L**). Collectively, we conclude that Snz-GFP is polarized to the cell periphery at ER-PM contact sites through association with the PM via phospholipid interactions that involve its C-terminal PX domain.

### Snz interacts with LDs via its C-Nexin domain

Close observation of Snz-GFP foci revealed they associate with pLDs that are intimately wrapped by PM invaginations (Figure 4D). Indeed, Snz-GFP foci often formed extended patches or “cups” that concentrated in the narrow spaces separating the PM and LD surface, suggesting that Snz-GFP foci may enrich between the PM and LD surface at sites of their close juxtaposition (Figure 5A). Similar “cups” have also been observed for proteins that enrich at ER-LD contact sites where LDs are budding from the ER surface, and Mdm1 forms such “cups” at ER-LD contact sites adjacent to the yeast vacuole, forming so-called ER-LD-vacuole *tri-organelle* contacts (Hariri et al., 2019).

**Figure 5:**
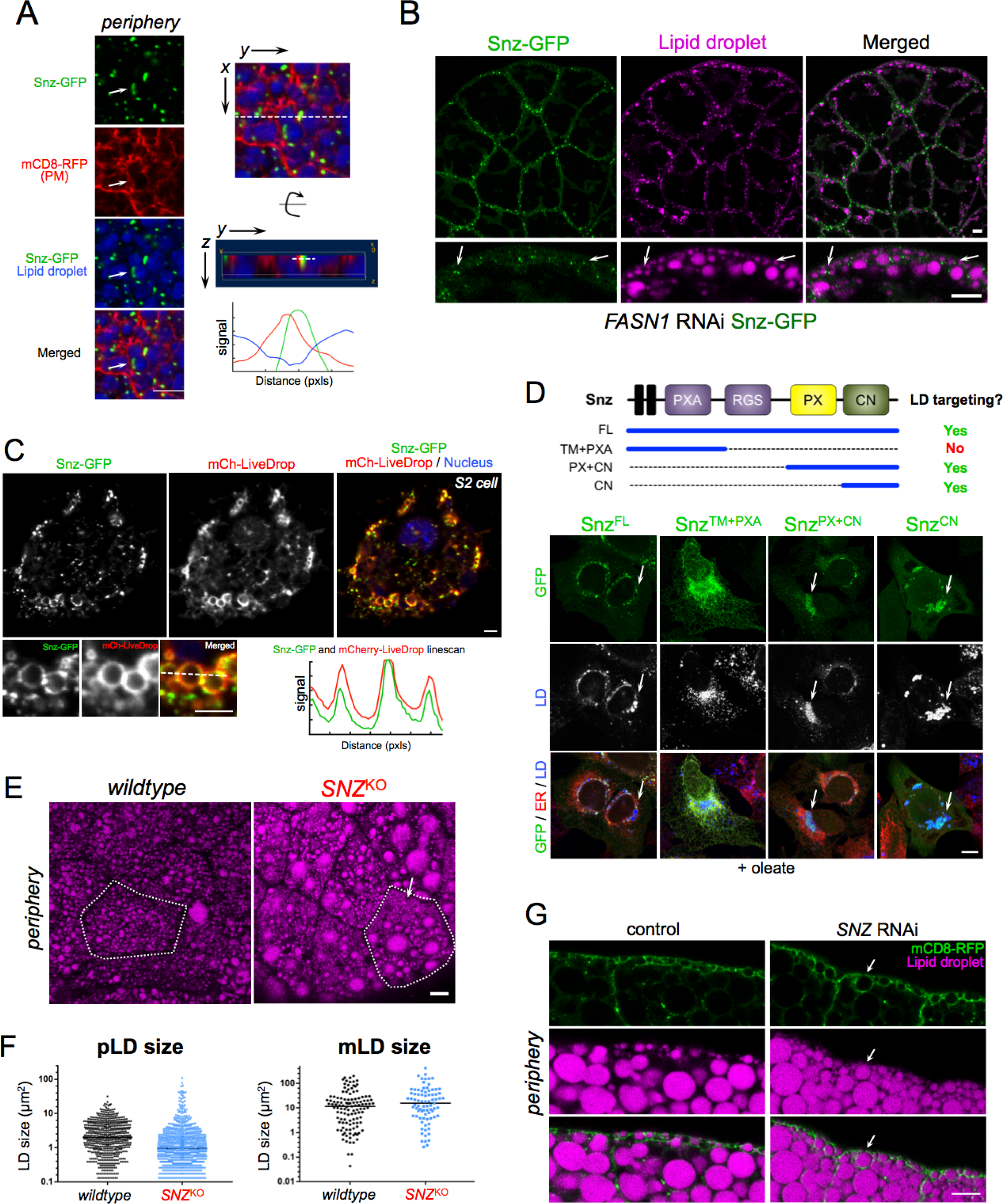
Snz interacts with LDs via its C-Nexin domain and loss of *SNZ* perturbs LD homeostasis in the FB. **A**) FB cell periphery expressing Snz-GFP (green) (*Dcg-Gal4>UAS-Snz-GFP*), PM marker mCD8-mCherry (red) (*Dcg-Gal4>mCD8-mCherry*), and stained for LDs (MDH, blue). Arrow indicates a Snz-GFP “cup” that is in close proximity to a pLD and PM structure. Linescan indicated with white line. This region is displayed both from the “top” x,y view as well as “side” y,z view. Scale bar 10µm. **B**) *FASN1*^RNAi^ FB (*Dcg-Gal4>UAS-FASN1*^*RNAi*^) expressing Snz-GFP (green) (*Dcg-Gal4>UAS-Snz-GFP*) and stained for LDs (MDH, magenta). Arrows indicate sites where Snz-GFP forms foci at the cell periphery that associate adjacent to pLDs. Scale bar 10µm. **C**) *Drosophila* S2 cell expressing Snz-GFP (green) and LD surface marker mCherry-LiveDrop (mCh-LiveDrop, red) and DAPI stained (blue). Linescan indicated with a white line. Scale bar 10µm. **D**) Mammalian *U2OS* cells treated overnight with oleate-conjugated BSA and expressing Snz-GFP fragments as denoted by the schematic at top of Panel. Snz-GFP fragments (green), ER marker is lumenal HSP90B1 (red) and LDs (MDH, blue, white in middle panel). Arrows indicate Snz-GFP targeting to LDs. Scale bar 10µm. **E**) FB from *SNZ*^KO^ larvae stained for LDs (MDH, magenta). A single cell is outlined in white. Arrow indicates a region of dense small pLDs. Scale bar 10µm. **F**) Dot plot of individual LD sizes in control *w*^*1118*^ and *SNZ*^KO^ larvae as represented in Panel E. **G**) FB side profiles of peripheries of control (*Dcg-Gal4* alone) and *SNZ*^RNAi^ (*Dcg-Gal4>UAS-SNZ*^*RNAi*^) larvae expressing mCD8-GFP PM marker (green) (*Dcg-Gal4>UAS-mCD8-mCherry*, that is pseudo-colored green here) and stained for LDs (MDH, magenta). Arrows indicate the absence of a clear pLD layer in *SNZ*^RNAi^ larvae. Scale bar 10µm.

To rule out the possibility that this Snz-LD association was not merely a consequence of the high LD density in FB cells, we imaged Snz-GFP in *FASN1*^*RNAi*^ FB tissue, which lacks mLDs but retains the pLD population in the cell periphery. Remarkably, Snz-GFP foci remained polarized to the cell periphery in *FASN1*^*RNAi*^ tissue, and associated with the surfaces of pLDs there (Figure 5B, **arrows**). To further investigate whether Snz-GFP was directly associating with LDs, we ectopically expressed Snz-GFP in *Drosophila* S2 cells co-expressing mCherry-LiveDrop, a known LD biomarker that accumulates on the LD monolayer surface (Wang et al., 2016). Following treatment with oleate to induce LD biogenesis, Snz-GFP co-localized with mCherry-LiveDrop on LD surfaces, suggesting Snz-GFP could interact directly with LD surfaces (Figure 5C).

To identify regions of Snz that mediate LD contact, we ectopically expressed Snz-GFP in mammalian *U2OS* cells. *U2OS* cells are an ideal model cell system to study protein targeting to LDs since they contain few LDs when cultured in standard DMEM media, but generate more LDs if the fatty acid oleate is added to the culture media. In the absence of oleate, GFP-tagged full-length Snz (Snz^FL^) localized to the ER tubular network, but was recruited to the surfaces of LDs following overnight incubation in media containing BSA-conjugated oleate (**SFigure 5A**). Next, we expressed GFP-tagged Snz fragments encoding either its N-terminal half containing the N-terminal transmembrane (TM) region and PXA domain (Snz^TM+PXA^), or the C-terminal half encoding the PX and C-Nexin domains (Snz^PX+CN^). In contrast, to Snz^FL^, Snz^TM+PXA^ failed to target to LDs in oleate-containing media, and instead localized throughout the ER network, co-localizing with the ER marker (Figure 5D, **SFigure 5B**). However, Snz^PX+CN^ targeted to LDs similar to Snz^FL^, indicating it was sufficient for LD targeting (Figure 5D, **SFigure 5B**). Similarly, GFP-tagged C-Nexin domain alone (Snz^CN^) also targeted to LDs following treatment with oleate. Collectively, we conclude that Snz can directly interact with LDs, and the C-Nexin domain is sufficient to mediate this LD targeting. This is similar to the Snz mammalian homolog SNX14 that also targets to LDs via its C-Nexin domain (Datta et al., 2019).

### Loss of *SNZ* perturbs pLD homeostasis

Given its ability to interact with LDs, we next investigated whether loss of Snz affected LD homeostasis in the FB. We generated a *Drosophila* line in which the entire *SNZ* open reading frame was deleted via CRISPR-Cas9 gene editing (denoted *SNZ*^KO^) (**SFigure 5C**). *SNZ*^KO^ larvae were healthy and manifested no obvious phenotypes under ambient conditions, and were able to pupate and grow into adult flies. However, imaging revealed dramatic changes to the morphology of the pLD population (Figure 5E). pLDs no longer appeared uniform, and pLD size distribution was altered. There was an increased number of very small pLDs as well as some very large pLDs, causing the pLD population to appear more varied in size compared to control *w*^1118^ samples (Figure 5E, F). In contrast, the mLD population exhibited only mild changes in LD size, suggesting loss of Snz primarily affected the pLD population (Figure 5F).

Next, we determined whether loss of *SNZ* affected PM wrapping around pLDs. We knocked-down *SNZ* using RNAi in mCD8-mCherry-expressing FB tissue (*Dcg-Gal4>UAS-SNZ^RNAi^; Dcg-Gal4>UAS-mCD8-mCherry*), and examined the cell periphery. Whereas control cells (*Dcg-Gal4* alone) exhibited normal PM wrapping around pLDs, *SNZ*^RNAi^ samples again exhibited more varied and irregular size distributions in the pLD population, recapitulating the pLD defects seen in *SNZ*^KO^ tissue (Figure 5G). Notably the pLD layer, usually distinct just below the cell surface, was now much less distinguishable from other LDs around it (Figure 5G, **arrow**). However, PM wrapping still occurred around pLDs, but this PM wrapping occurred around both small and large pLDs, causing the PM to appear “bubbled” and irregular at cell edges (Figure 5G, **SFigure 5D arrows**). Collectively, this suggests that loss of Snz perturbs pLD morphology, but PM wrapping around pLDs can still occur in the absence of Snz.

Earlier we had noted that the morphology of the pLD population drastically changed during larval fasting, suggesting the TAG in pLDs was mobilized during fasting (Figure 2A-C). Since *SNZ*^KO^ larvae displayed alterations in pLD morphology, we next investigated whether *SNZ*^KO^ larvae would exhibit changes in fasting-induced mobilization of TAG. Control *(w^1118^)* and *SNZ*^KO^ larvae were fasted for 24 hours and their TAG levels monitored at 12 and 24 hours post-fasting by thin layer chromatography (TLC). Intriguingly, prior to fasting *SNZ*^KO^ larvae exhibited slightly elevated TAG levels compared to control larvae, suggesting some alteration in LD homeostasis and TAG storage (**SFigure 5E**). However, *SNZ*^KO^ larvae exhibited a more significant loss of TAG following 24 hours of fasting compared to *wildtype* (**SFigure 5E**). There was also a corresponding elevation of free fatty acids (FFAs) in *SNZ*^KO^ larvae, suggesting alterations in some aspect of fasting-induced TAG mobilization or LD homeostasis (**SFigure 5F**). Altogether, we conclude that *SNZ*^KO^ larvae manifest three phenotypes associated with LD and TAG homeostasis: 1) alterations in pLD morphology in fed conditions, 2) changes in TAG and FFA levels in fed conditions, and 3) alterations in fasting-induced TAG mobilization.

### Snz over-expression promotes TAG storage in the FB

Given its ability to target to LDs and its requirement for regular pLD morphology, we hypothesized that Snz regulated some aspect of TAG storage in LDs within the FB. To interrogate this, we monitored TAG levels in control (*Dcg-Gal4* driver alone) and transgenic *Drosophila* over-expressing Snz specifically in the FB (*Dcg-Gal4>UAS-Snz*, denoted as Snz-Transgenic, Snz-Tg). Intriguingly, both Snz-Tg larvae and 7-10 day old Snz-Tg flies contained significantly elevated TAG stores (Figure 6A,B). In line with this, Snz-Tg larvae exhibited significantly larger pLDs compared to controls (Figure 6C,D, **SFigure 6A,B**).

**Figure 6:**
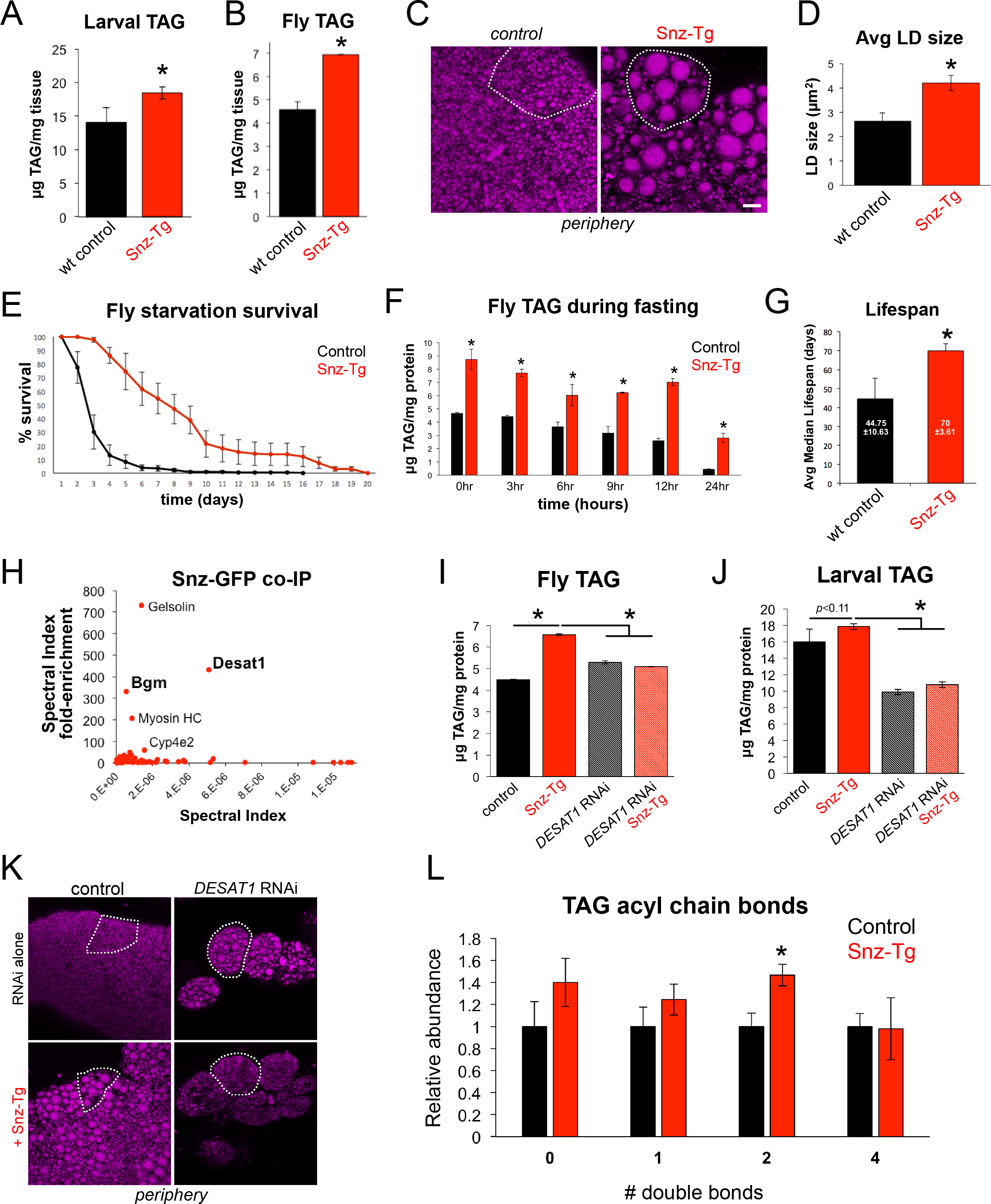
Tissue-specific Snz over-expression promotes TAG storage and organismal homeostasis in a DESAT1-dependent manner. **A**) Total larval TAG levels in control (*Dcg-Gal4* alone) and Snz transgenic (Tg) larvae with FB-specific over-expression of Snz (*Dcg-Gal4>UAS-Snz*). * indicates p<0.05. **B**) Total adult fly TAG from control (*Dcg-Gal4*) and Snz-Tg flies. * indicates p<0.05. **C**) FB tissue peripheries of control (*Dcg-Gal4*) or Snz-Tg and stained for LDs (MDH, magenta). Single cells are outlined in white. Scale bar 10µm. **D**) Average LD size in the periphery of control and Snz-Tg tissue as in Panel C. * indicates p<0.05. **E**) Starvation survival assay using 7-10 day old control (*Dcg-Gal4*) or Snz-Tg flies over-expressing Snz in the FB. **F**) Total fly TAG levels of control and Snz-Tg flies at time intervals during a starvation assay as in Panel E. * indicates p<0.05. **G**) Median lifespan of control (*Dcg-Gal4*) and Snz-Tg flies reared on a normal diet. * indicates p<0.05. **H**) Fold-enrichment plot for mass spectrometry-based proteomics of Snz-GFP (*Dcg-Gal4>UAS-Snz-GFP*) and soluble GFP (*Dcg-Gal4>UAS-GFP*) immuno-precipitates from larval tissue. **I**) Total fly TAG in control (*Dcg-Gal4*), Snz-Tg, FB-specific *DESAT1*^RNAi^ (*Dcg-Gal4>UAS-DESAT1*^*RNAi*^), and *DESAT1*^RNAi^ + Snz-Tg (*Dcg-Gal4>UAS-DESAT1^RNAi^; Dcg-Gal4>UAS-Snz-GFP*). * indicates p<0.05. **J**) Total larval TAG for 7-10 day old flies for samples as in Panel I. * indicates p<0.05. **K**) FB peripheries stained for LDs (MDH, magenta) in control (*Dcg-Gal4*) or *DESAT1*^RNAi^ samples. Single cells are outlined in white. Scale bar 10µm. **L**) Relative abundances of # of double bonds in TAG-associated fatty acyl chains from total TAG in control (*Dcg-Gal4*) and Snz-Tg (*Dcg-Gal4>UAS-Snz*) larvae. * indicates p<0.05.

Since TAG provides nutrients during long-term fasting, we determined whether the elevated TAG stores in Snz-Tg flies could promote their survival during sustained food deprivation. We monitored 7-10 day old Snz-Tg and control flies during persistent starvation where they were supplied with water but no food. Intriguingly, Snz-Tg flies survived significantly longer than controls during persistent nutrient starvation (Figure 6E). We also monitored Snz-Tg and control TAG levels at several time-points throughout the first 24 hours of starvation. Both Snz-Tg and control flies gradually mobilized TAG, but Snz-Tg flies exhibited consistently higher TAG levels compared to controls (Figure 6F, **SFigure 6C**). Collectively, we conclude that FB-specific over-expression of Snz is sufficient to elevate TAG stores which can be harvested during long-term fasting.

### Tissue-specific Snz over-expression promotes organismal lifespan and homeostasis

In addition to their role as a nutrient reservoir, TAG stores can also promote general organismal homeostasis by serving as a metabolic buffer for changes in dietary nutrient availability and flux. Consequently, elevated TAG stores have been shown to extend organismal lifespan in *Drosophila* and other model organisms (Bjedov et al., 2010) (Palgunow et al., 2012) (Handee et al., 2016). Since Snz-Tg flies exhibited elevated TAG stores, we monitored a population of Snz-Tg (*Dcg-Gal4>UAS-Snz*) and control flies fed a normal diet over the course of their entire life. Strikingly, Snz-Tg populations exhibited a significantly longer median lifespan (~70 days ±4) compared to controls (~45 days ±11), indicating over-expression of Snz in the FB was sufficient to extend *Drosophila* lifespan (Figure 6G).

Animals fed excess nutrients can exhibit elevated sugar and lipid levels in their blood. In *Drosophila*, FB TAG stores have been proposed to serve as sinks for these excess nutrients which can otherwise contribute to hyperglycemia and insulin resistance similar to type 2 diabetes (T2D) in mammals. In line with this, perturbation of TAG synthesis in the FB hyper-sensitizes *Drosophila* to the caloric overload caused by a high-sugar diet (HSD) (Musselman et al., 2011) (Musselman et al., 2013). Given the ability of Snz to promote TAG storage in the FB, we interrogated whether Snz-Tg flies were resistant to the pathological effects of a chronic HSD. We reared 7-10 day old Snz-Tg (*Dcg-Gal4>UAS-Snz*) and control (*Dcg-Gal4* alone) flies on a chronic diet of normal food (5% sugars) or a HSD containing 35% sugars, and monitored their long-term survival. As expected, control flies fed the HSD diet exhibited morbidity and decreased survival compared to the normal diet (**SFigure 6D**). In contrast, Snz-Tg flies survived similarly on both diet regimens, and appeared healthy and generally unaffected by the HSD (**SFigure 6D**). Collectively, we conclude that Snz over-expression in the FB is sufficient to promote TAG storage, and these elevated TAG stores can promote survival during starvation as well as contribute to organismal homeostasis in animals fed a chronic HSD.

### Snz functionally interacts with fatty acid desaturase DESAT1

To begin to characterize the molecular function of Snz, we conducted co-immunoprecipitation-based mass spectrometry to identify potential Snz interacting partners. We expressed either Snz-GFP (*Dcg-Gal4>UAS-Snz-GFP*) or soluble GFP (*Dcg-Gal4>UAS-GFP*) in the larval FB and co-purified both with their associated proteins using an anti-GFP affinity resin. Mass spectrometry-based proteomics revealed several proteins that preferentially co-purified with Snz-GFP, including actin-associated proteins gelsolin and myosin heavy chain (Myosin HC) (Figure 6H, **SFigure 6E**). We also detected the fatty acyl-CoA ligase Bubblegum (BGM) and fatty acid (FA) desaturase DESAT1, which are involved in the activation and desaturation of FAs, respectively (Min and Benzer, 1999) (Pei et al., 2003) (Musselman et al., 2013) (Figure 6H). DESAT1 is a member of the stearoyl-CoA desaturase family, and acts on long chain saturated FAs to convert them to unsaturated FAs prior to their incorporation into TAG (**SFigure 6H**). Notably, DESAT1 expression is up-regulated in flies fed a HSD, and its tissue-specific knockdown in the FB reduces TAG stores and hyper-sensitizes flies to HSD-associated toxicity (Musselman et al., 2013).

Given its importance in FA metabolism and TAG synthesis in the FB, we investigated whether the elevated TAG stores observed in Snz-Tg animals required DESAT1. We measured TAG and FFA levels in control (*Dcg-Gal4* alone) and Snz-Tg flies in either the presence or absence of *DESAT1* in the FB (*Dcg-Gal4>UAS-DESAT1^RNAi^*). Strikingly, knockdown of DESAT1 in Snz-Tg adult flies suppressed their TAG accumulation, and Snz-Tg flies now contained TAG levels similar to controls (Figure 6I). In line with this, *DESAT1*^RNAi^ flies over-expressing Snz (Snz-Tg) also contained elevated FFAs, suggesting a block in FA incorporation into TAG (**SFigure 6F**). Next, we knocked-down *DESAT1* in the larval FB, and examined their TAG and FFAs. Consistent with previous reports, *DESAT1*^*RNAi*^ larvae were lipodystrophic and their FB cells appeared deflated and poorly connected to one another (Figure 6K) (Musselman et al., 2013). Snz over-expression in the *DESAT1*^*RNAi*^ larvae failed to rescue this tissue lipodystrophy, suggesting DESAT1 was required for Snz-driven TAG accumulation in larvae (Figure 6K). Similarly, TAG levels in *DESAT1*^*RNAi*^ + Snz-Tg larvae mirrored controls, further indicating that DESAT1 is required for Snz-driven TAG accumulation (Figure 6J). *DESAT1*^*RNAi*^ + Snz-Tg larvae also contained elevated FFAs, consistent with a defect in FA incorporation into TAG (**SFigure 6G**). Thus we conclude that DESAT1 is required for Snz-driven TAG accumulation in both flies and larvae.

### FB-specific Snz over-expression alters the TAG acyl chain lipidome

Since Snz interacted with enzymes involved in FA metabolism, we also investigated whether Snz-Tg larvae had alterations in their TAG-conjugated acyl chains. We conducted mass spectrometry-based lipidomic profiling of TAG-derived FAs from Snz-Tg and control larvae. Quantitative lipidomic profiling revealed changes in the relative abundances of several TAG-derived FAs in Snz-Tg larvae. In particular, Snz-Tg larvae displayed a significant increase in the relative abundance of acyl chains with two double bonds (Figure 6L, **SFigure 6I**). Lipid profiling revealed that this was primarily due to an increased abundance of the essential polyunsaturated FA (PUFA) linoleic acid (18:2) in the TAG pool of Snz-Tg larvae (**SFigure 6I**).

Collectively we conclude that Snz functionally interacts with the FA desaturase DESAT1 and requires DESAT1 to stimulate TAG storage in the FB. Tissue-specific over-expression of Snz in the FB promotes TAG accumulation but also alters the FA profile of TAG-associated acyl chains, and leads to an increase in the relative abundance of PUFAs with two double bonds in TAG stores.

## Discussion

Professional fat-storing cells must organize their fat reserves to balance long-term storage with the ability to efficiently mobilize lipids during energetic crises like starvation or metamorphosis. How this organization is achieved is unknown but presents significant spatial and metabolic challenges for the cell. Here, we report that *Drosophila* FB adipocytes contain functionally distinct LD populations that are spatially segregated in the cell cytoplasm (Figure 7A). A pLD population is maintained adjacent to the cell surface and makes intimate contact with the PM. pLD size and abundance are altered in response to fasting, suggesting pLDs are mobilized to provide circulating lipids for other larval tissues. Consistent with this, loss of lipophorin (Lpp) particles by *ApoLpp*^RNAi^ impacts pLD abundance and morphology, suggesting pLD maintenance requires some aspect of Lpp lipid trafficking. FB cells also contain a larger mLD population in the cell mid-plane region that is unaffected by loss of Lpp, but is drastically affected by loss of FASN1-mediated *de novo* lipogenesis in the FB. Remarkably, pLDs were still observed docked on the inner surface of the PM in *FASN1*^*RNAi*^ FB tissue, further suggesting that pLDs contain lipids derived from extracellular sources that may be delivered into the FB via Lpp-dependent trafficking. We also find that pLDs and mLDs are differentially dependent on perilipins, with mLDs relying on LSD1 for their morphology whereas pLDs require LSD2. Finally, we identify Snz as a LD-associated protein that is required for proper LD homeostasis in the FB. Snz localizes to ER-PM contacts in the FB cell periphery, and its over-expression increases TAG storage, consistent with a model whether Snz regulates ER-PM inter-organelle crosstalk that promotes lipid storage in LDs. In line with this, Snz functionally interacts with the ER-resident FA desaturase DESAT1, which is required for Snz-driven TAG accumulation.

**Figure 7:**
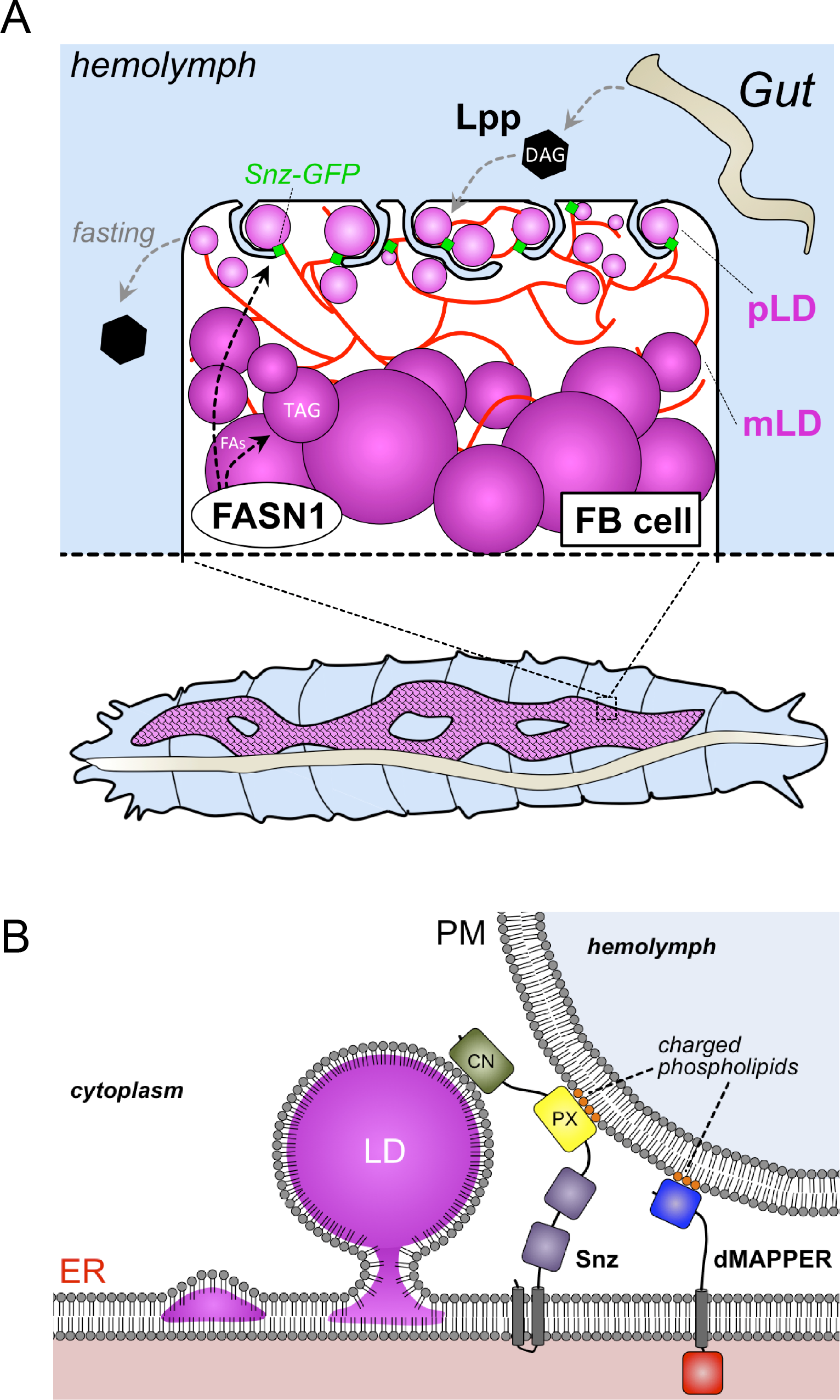
Working models for Snz in Drosophila FB cell LD homeostasis. **A**) Working model depicting spatially distinct LD populations in *Drosophila* FB cells. **B**) Topological cartoon of Snz interacting with the ER, PM, and LDs in *Drosophila* FB cell. The DMAPPER ER-PM tether is also depicted. Scale bar 10µm.

LDs have long been observed to be tethered to other organelles such as mitochondria and peroxisomes, and this impacts their sub-cellular distribution as well as their ability to exchange lipids with these organelles (Olzmann and Carvalho, 2019). Recent studies have identified specific proteins that bind to the surfaces of LDs, and mediate their attachment to other cellular organelles. Among these, yeast Mdm1 directly binds to nascent LDs, and promotes their attachment to the yeast vacuole. This Mdm1-vacuole interaction is critical for defining this LD positioning, as replacement of Mdm1’s vacuole-binding PX domain with a PM-binding domain re-localizes Mdm1 to ER-PM contact sites and causes LDs to bud instead from the cortical ER adjacent to the PM (Hanaa Hariri, 2019). In addition to its role as an organelle tether, Mdm1 also positively regulates LD biogenesis by recruiting the fatty acyl-CoA ligase Faa1 to the ER surface, where it induces the incorporation of FAs into TAG in the LD (Hanaa Hariri, 2019). Here, we find that Snz may function similarly to Mdm1 in both the spatial positioning of LDs, as well as promoting LD biogenesis. We find that Snz localizes to the FB cell periphery and co-localizes with the ER-PM contact site biomarker dMAPPER, suggesting it enriches at regions of close contact between the ER and PM and potentially functions as an inter-organelle tether (Figure 7B). Consistent with this, the Snz PX domain binds to liposomes containing phospholipids normally enriched on the PM, and exhibits a non-canonical lipid binding surface that mediates electrostatic interactions with phospholipids that would be present on the PM (Chandra M, 2019). This suggests a model where Snz functions in some aspect of functional coupling between the ER and PM, and potentially assists in lipid uptake from the hemolymph. Snz may also help to localize ER-resident lipid processing enzymes such as DESAT1 to the cell periphery, creating a localized pool of FA processing enzymes in the peripheral ER network which can efficiently process lipids prior to their incorporation into TAG. Consistent with this, Snz over-expression in the FB promotes TAG storage, and Snz can directly associate with LDs.

Once thought to be pathogenic, adipocyte fat stores have more recently been proposed to act as metabolic buffers that protect against caloric overload and serve as sinks to reduce circulating lipids and sugars. As such, factors that enhance adipocyte fat storage may be protective against insulin insensitivity, organismal lipotoxicity, and other T2D-like pathologies (Musselman et al., 2013) (Musselman et al., 2011) (Listenberger et al., 2003). Our data are consistent with this model, and indicate that Snz up-regulation promotes TAG storage in the FB through a DESAT1-dependent pathway. These elevated TAG stores not only prplong organismal survival during sustained fasting, but also promote organismal homeostasis that extends *Drosophila* lifespan, as well as buffers the pathological effects of chronic HSD.

Collectively, our data support a model where Snz-associated pLDs provide a spatially-compartmentalized sink for lipids derived from the extracellular hemolymph. This pLD population may serve multiple functions. It could allow FB cells to quickly and efficiently process and store incoming Lpp-derived lipids from the hemolymph, thus avoiding their potentially lipotoxic accumulation in the cytoplasm. This would promote general cellular homeostasis and could minimize FA lipotoxicity during elevated lipid uptake. In addition, the high surface-to-volume size ratio of small pLDs may promote their efficient mobilization by cytoplasmic lipases during fasting. Since they are near the surface, pLD mobilization could also allow liberated FAs to be efficiently transferred to Lpp particles docked on the FB surface, where they can be subsequently trafficked to other organs. Snz has clear mammalian homologs including SNX25 which also bind PM-associated phospholipids (Chandra and Collins, 2018). Whether SNX25 or other Snz homologs are also able to interact with LDs in distinct regions of the mammalian cell interior is unclear, but will be the focus of future studies.

## Supporting information

SFigures

SMovie_1

## Acknowledgements

The authors would like to thank Drs. Sandra Schmid and Joel Goodman for help and comments in preparing this manuscript. We also greatly thank Dr. Jen Liou for the MAPPER construct. We would also like to thank Drs. Kate Luby-Phelps and Anza Dareshouri for assistance with the TEM. We also thank Helmut Kramer, Drew Stenesen, Andrew Moehlman and other members of the Kramer and Graff labs for helpful comments and suggestions. This work is supported by grants from the Welch Foundation (I-1873), the Searle Foundation (SSP-2016-1482), the NIH NIGMS (GM119768), AFAR (A15198), and the UT Southwestern Endowed Scholars Program. BMC is supported by an Australian National Health and Medical Research Council (NHMRC) Senior Research Fellowship (APP1136021). JGM is supported in part by NIH 2 PO1 HL20948-33.

## Author Contributions

WMH and RU developed the overall concepts and experiments. SJ, JB, MFB, HH, AV, and SF assisted with fly experiments. HH conducted the Snz-GFP immuno-precipitation and proteomics experiments. SR assisted in data quantification. MC and BC performed the Snz PX structure modeling and liposome binding assays.

## Materials and Methods

### Fly culture and strains

Flies were reared and maintained on standard fly food containing cornmeal, yeast, molasses (~5% sugars), and agar. High-sugar diet (HSD) media containing 35% sugars was prepared by adding additional sucrose (30 mg/ml). The *w*^*1118*^ common laboratory control strain was obtained from WellGenetics (Taiwan) where the *SNZ* knockout (KO) null strain was also generated (details on KO generation below). The Dcg*-*Gal4 fat body (FB)-specific driver, previously described in (Suh et al., 2008), and mCD8-GFP strains were provided by Jonathan M. Graff (UTSW) and Helmut Kramer (UTSW), respectively. Constructs for UAS-SNZ FL-GFP, UAS-SNZ FL (untagged), and UAS-SNZ(ΔC)-EGFP strains were generated in the Henne Lab and flies transformed by Rainbow Transgenic Flies, Inc. (Camarillo, CA). All other TRiP RNAi stocks and protein/organelle marker strains used were obtained from the Bloomington Stock Center (Bloomington, IN). All crosses involving Gal4 and UAS lines were cultured at 25°C to maintain uniform transgene expression. Unless otherwise indicated, all RNAi knockdowns were conducted using the Dcg-Gal4 UAS driver system.

### SNZ CRISPR mutant generation

WellGenetics engineered and generated the *SNZ*^KO^ mutant. Briefly, a gRNA located in the 5’ UTR (GAAGGAGAGGTCGTCGACAA) and another one located 83bp upstream of the stop codon (GTAATGCGATTGGTGGTGCG) were cloned into pDCC6 vectors. *w*^*1118*^ flies were injected with both gRNA expressing plasmids along with a recombination cassette encoding a hsp70::w marker with homology arms corresponding to the 5’ and 3’ ends of the *SNZ* gene. Red eyed progenies were screened by PCR and sequencing for insertion of the hsp70::w cassette into the *SNZ* locus. One such line showed proper insertion of the cassette into the *SNZ* locus and resulted into a deletion starting at 188 bp upstream of *SNZ* ATG-start codon up to 83 bp after the stop codon, causing a 4.5 kb deletion completely removing *SNZ* coding sequence.

### Molecular cloning and transgenic constructs

All constructs were amplified from a home-made *Drosophila* cDNA library, and cloned into either the pUASt vector for *Drosophila* expression or N2-pEGFP for mammalian U2OS cell expression. Snz FL (residues 1-1117) or Snz(ΔC) (residues 1-390) were fused to a C-terminal EGFP and inserted via EcoRI/XbaI cut sites. The mCherry-LiveDrop was cloned from *Drosophila* cDNA based on the sequence of *D.m.* GPAT4 from (Wang et al., 2016). The UAS-mCD8-GFP, UAS-mCD8-mCherry, and UAS-Sec61-TdTomato marker lines were obtained from the Bloomington Stock Center.

### S2 Cell Imaging

5 × 10^5^ S2 cells were plated per well in 6-well plates in Schneider’s complete media. On the next day, cells were transfected with 0.5µg of Met-GAL4 plasmid, 1 µg of SNZ(FL)-GFP expression plasmid and 1µg of mCherry-LiveDrop expression plasmid using TransIT^Tm^-Insect transfection reagent following manufacturer’s instructions. Twenty-four hours later, transgene expression was induced by the addition of 0.5mM copper sulfate and cells were incubated for 16 hours. Cells were removed by pipetting and transferred into 4 independent wells of a 24 well plates containing a #1.5 coverglass previously coated with a solution of 0.5mM solution of Concanavalin A. Cells were left to adhere for 1 hour and then were incubated with either the vehicle (EtOH) or 1mM Oleate in Schneider media for 45 minutes. Cells were then process for immunofluorescence analysis as described for U2OS cells and imaged on an upright LSM880 confocal microscope with a 40×/1.4NA Plan Apo oil objective.

### Confocal fluorescence microscopy

Mid-third instar feeding larvae were gently removed from inside the food media using a paintbrush, and rinsed in water to remove food particles. Fat bodies (FBs) were dissected from larvae in PBS using Dumont #5 forceps (Electron Microscopy Sciences). All FBs were fixed in 4% paraformaldehyde for 30 minutes at room temperature and rinsed briefly in PBS, prior to staining with organelle labeling dyes or mounting directly in SlowFade Gold antifade reagent with DAPI (Invitrogen) if FBs transgenically expressed fluorescent-tagged proteins. Prepared slides were refrigerated overnight at 4°C before imaging using Zeiss LSM880 inverted laser scanning confocal microscope. Most FBs were imaged using a 40X oil immersion objective, as 1-µm z-stack sections when imaging from periphery to mid-plane (ie. peri-nuclear region) of cells, or 2.5-4 µm sections when imaging from apical to basal surface of the ~40 µm thick monolayer.

### Lipidomic Profiling

#### Chemicals and Materials

All solvents were either HPLC or LC/MS grade and purchased from Sigma-Aldrich (St Louis, MO, USA). SPLASH LipidoMix™ standards, a mixture of deuterated lipids from major lipid classes found in biological samples, were purchased from Avanti Polar Lipids (Alabaster, AL, USA). All lipid extractions were performed in 16×100mm glass tubes with PTFE-lined caps (Fisher Scientific, Pittsburg, PA, USA). Glass Pasteur pipettes and solvent-resistant plasticware pipette tips (Mettler-Toledo, Columbus, OH, USA) were used to minimize leaching of polymers and plasticizers.

#### Sample homogenization

Approximately two fruit flyes were transferred to a 2.0-mL pre-filled Bead Ruptor tube (2.8mm ceramic beads, Omni International, Kennesaw, GA, USA), and 1mL of methanol/dichloromethane (1:2, v/v) was added. Tissue was homogenized with a Bead Ruptor (Omni International) for 50 seconds (5.5 mps, 3 cycles, 10 sec/cycle, 5 s dwell time). The homogenates were transferred to glass tubes for Liquid Liquid Lipid extraction (LLE).

#### Liquid-Liquid Lipid Extractions

LLE were performed at room temperature (including centrifugation) to maintain consistent solubility and phase separation. The modified Bligh/Dyer extraction began by adding 1mL each of methanol, dichloromethane, and water to a glass tube containing the sample (see Sample Preparation). The mixture was vortexed and centrifuged at 2671*×g* for 5 minutes, resulting in two distinct liquid phases. The organic phase (bottom phase) was collected to a fresh glass tube with a Pasteur pipette and dried under N_2_. The dried extracts were reconstituted in 600µL of dichloromethane/methanol/isopropanol (2:1:1, v/v/v) containing 8mM ammonium fluoride (NH_4_F) and 33 µL/mL of 3:50 diluted SPLASH LipidoMix™ internal standard.

#### Lipids Profiling by Direct-Infusion MS/MS^all^ workflow

Lipid extracts were infused into a SCIEX quadrupole time-of-flight (QTOF) TripleTOF 6600+ mass spectrometer (Framingham, MA, USA) via a custom configured LEAP InfusePAL HTS-xt autosampler (Morrisville, NC, USA). Electrospray ionization (ESI) source parameters were, GS1 25, GS2 55, curtain gas (Cur) 20, source temperature 300 °C and ion spray voltage 5500V and −4500V in positive and negative ionization mode, respectively. GS1 and 2 were zero-grade air while Cur and CAD gas was nitrogen. Optimal declustering potential and collision energy settings were 120V and 40eV for positive ionization mode and −90V and −50eV for negative ionization mode. Samples were infused for 3 minutes at a flow rate of 10µL/min. MS/MS^ALL^ analysis was performed by collecting product-ion spectra at each unit mass from 200-1200 Da (sequentially adjusted for mass-defect of hydrogen) with an accumulation time of 0.3 seconds per mass. Analyst^®^ TF 1.7.1 software (SCIEX) was used for TOF MS and MS/MS^ALL^ data acquisition. Chronos XT^®^ software from Axel Semrau (Srockhovel, Germany) was used to control the InfusePAL system. Data analysis was performed by MarkerView (SCIEX) peak-picking algorithm to generate precursor/fragment pairs from MS/MS^ALL^ spectra (mass tolerance window of 0.3 Da and an intensity threshold of 30), followed by species identification using LipPy, an in-house script. This script provides instrument quality control information, isotopic peak corrections, lipid species identification, data normalization, and basic statistics.

**SFigure 1: Peripheral LDs (pLDs) associate with the PM**. **A**) Cartoon schematic of *Drosophila* larval fat body (FB) tissue with intra-cellular LDs (magenta). PM marker mCD8-GFP is denoted in green. **B**) FB tissue expressing a GFP-tagged PtdIns(4,5)P_2_ binding biomarker PLCδ-2XPH domain (green, pseudo-color) (*Dcg-Gal4>UAS-mCherry-PLCδ-2XPH*) and stained for LDs (MDH, magenta). Scale bar 10µm. **C**) FB tissue from 7-10 day old flies expressing mCD8-GFP PM marker (green) (*Dcg-Gal4>UAS-mCD8-GFP*) and stained for LDs (MDH, magenta). Scale bar 10µm. **D**) TEM micrographs of L3 larval FB tissue showing FB cell periphery with pLDs and PM-associated invaginations. Black arrows denote PM sites of invagination. Red arrows denote ER tubular networks in the cell periphery. Scale bar 1µm. **E**) TEM of different field of view FB tissue as in Panel D. Arrows as in Panel D. Scale bar 1µm.

**SFigure 2: pLDs and mLDs are differentially affected by loss of ApoLpp and FASN1. A**) mLD size distributions in FB tissue from fed non-wandering L3 larvae, 24 hr fasted larvae, and larvae fed for 24 hr on a high-sugar diet (HSD) containing 35% sugars. * indicates p<0.05. **B**) mLD densities in fed larvae, 24 hr fasted larvae, and 24 HSD-diet fed larvae. * indicates p<0.05. **C**) FB tissue from control (*Dcg-Gal4* alone) and *ApoLpp*^RNAi^-treated tissue (*Dcg-Gal4>UAS-ApoLpp^RNAi^*) stained for LDs (MDH, magenta). Arrows indicate pLDs. Scale bar 10µm. **D**) Average mLD size for control (*Dcg-Gal4* alone) and *ApoLpp^RNAi^*-treated larvae. **E**) Average mLD density for control (*Dcg-Gal4* alone) and *ApoLpp*^*RNAi*^-treated larvae. **F**) FB tissue from control (*Dcg-Gal4* alone) and *ApoLpp*^*RNAi*^-treated larvae. FB tissue is expressing mCD8-GFP PM marker (green) (*Dcg-Gal4>UAS-mCD8-GFP*) and stained for LDs (MDH, magenta). Left panels are side-views of the tissue edge, and right panels are the top profile of a cell surface. Scale bar 10µm. **G**) Total larval TAG for control (*Dcg-Gal4* alone) of *ApoLpp*^*RNAi*^-treated treated larvae. **H**) Total larval DAG for control (*Dcg-Gal4* alone) and *ApoLpp*^*RNAi*^-treated larvae. * indicates p<0.05. **I**) Total larval TAG for control (*Dcg-Gal4* alone) and *FASN1^RNAi^-*treated larvae (*Dcg-Gal4>UAS-FASN1^RNAi^)*. * indicates p<0.05. **J**) Total larval DAG for control (*Dcg-Gal4* alone) and *FASN1^RNAi^-*treated larvae as in Panel I.

**SFigure 3: pLDs are perturbed by loss of β-spectrin and LSD2. A**) Average mLD size for control (*Dcg-Gal4* alone) and β*-spectrin*^RNAi^ knockdown (*Dcg-Gal4>UAS-β-spectrin^RNAi^*) in the FB. * indicates p<0.05. **B**) Average mLD density for control (*Dcg-Gal4* alone) and β*-spectrin*^RNAi^ knockdown in the FB. * indicates p<0.05. **C**) FB tissue stained for LDs (MDH, magenta) from control (*Dcg-Gal4* alone) and *LSD2*^RNAi^ in the FB (*Dcg-Gal4>UAS-LSD2^RNAi^*). Individual cells are outlined in white. Arrows indicate pLDs. Scale bar 10µm. **D**) Average pLD size for control (*Dcg-Gal4* alone) and *LSD2^RNAi^* knockdown (*Dcg-Gal4>UAS-LSD2^RNAi^*) in the FB. * indicates p<0.05. **E**) Average pLD densty for control (*Dcg-Gal4* alone) and *LSD2^RNAi^* knockdown in the FB. * indicates p<0.05. **F**) Average mLD size for control (*Dcg-Gal4* alone) and *LSD2*^*RNAi*^ knockdown (*Dcg-Gal4>UAS-LSD2*^*RNAi*^) in the FB. Samples are not significantly (n.s.) different. **G**) Average mLD density for control (*Dcg-Gal4* alone) and *LSD2*^*RNAi*^ knockdown (*Dcg-Gal4>UAS-LSD2*^*RNAi*^) in the FB. Samples are not significantly (n.s.) different.

**SFigure 4: Snz-GFP is polarized to the cell periphery**. **A**) Relative expression of endogenous SNZ as determined by qRT-PCR for endogenous *SNZ* cDNA from L3 larvae and 7-10 day old *Drosophila* flies. **B**) Immuno-fluorescence stain of Snz-GFP (anti-GFP, green) and ER marker (anti: Calnexin 99a, red) in larval FB tissue. White box depicts the inset images. Scale bar 10µm. **C**) Gray-scale z-stack images of FB tissue expressing Snz-GFP (black foci in image) in sections through FB tissue. Z-section number through tissue is shown in upper-right corner. Step-sizes of sections were 5 µm through tissue. Total membrane stain is Cell Mask (red). Nuclei is in blue from DAPI stain. **D**) Quantification of total Snz-GFP foci per section of FB tissue as shown in Panel C. **E**) Quantification of % Snz-GFP foci by depth into FB tissue using 1µm z-stack sections for 15 sections into FB tissue. ~65% of Snz-GFP foci detected were within the first 5 sections of tissue from surface. **F**) Snz-GFP (green) expressing larval FB tissue (*Dcg-Gal4>UAS-Snz-GFP*) stained for LDs (MDH, magenta). Arrows indicate Snz-GFP foci at cell periphery in peripheral (left) or mid-plane (right) sections of FB tissue. Scale bar 10µm. Cartoon below depicts Snz-GFP foci in same z-section plane with the pLDs. **G**) (*left*) Side-profile of periphery of Snz-GFP (green) expressing FB tissue (*Dcg-Gal4>UAS-Snz-GFP*) also expressing ER marker Sec61-TdTomato (red) (*Dcg-Gal4>UAS-Sec61-TdTomato*) and stained for LDs (MDH, blue). Arrows depict Snz-GFP foci at ER-positive regions. (*right*) Side-profile of periphery of Snz-GFP (green) expressing FB tissue (*Dcg-Gal4>UAS-Snz-GFP*) also expressing PM marker mCD8-mCherry (red) (*Dcg-Gal4>UAS-mCD8-mCherry*) and stained for LDs (MDH, blue). White lines in the images shows regions depicted in the linescans on far right. Scale bar 10µm. **H**) Peripheries of FB cells expressing either full length Snz-GFP or Snz^ΔC^-GFP (green). Cartoons depict potential topological localization of these constructs. Scale bar 10µm. **I**) Molecular ribbon model (*left*) and surface electrostatic charge (*right*) of *Drosophila* Snz PX domain modeled on NMR structure of SNX25 PX domain (PDB: 5WOE). Residues forming the canonical PtdIns(3)P binding pocket are labeled in magenta. Residues forming the non-canonical positively-charged patch are in blue. Below these structures is a primary sequence comparison of SNX13, SNX25, and SNZ PX domains region showing residues involved in canonical (red) and non-canonical (blue) lipid-binding surfaces. Secondary structures also depicted. **J**) Liposome co-sedimentation assays of *wildtype* or mutant Snz PX domains. PC: phosphatidylpholine, PE: phosphatidylethanolamine liposomes. **K**) Quantification of Pellet (P) to Supernatent (S) ratios for experiments depicted in Panel J. **L**) Top views of larval FB tissue from control (*Dcg-Gal4 alone*) or dPIP5K RNAi –treated larvae. Tissue is expressing PM marker mCD8-GFP (*Dcg-Gal4>UAS-mCD8-GFP*) and stained for LDs (MDH, magenta). PM wrapping around pLDs still occurs.

**SFigure 5: Snz interacts with LDs via its C-Nexin domain and its loss perturbs LD homeostasis. A**) *U2OS* cells expressing full length Snz-GFP and cultured in DMEM media either without (*left*) or with (*right*) BSA-conjugated oleate. Lipid droplets (MDH, magenta) also stained. White boxes indicate insets at bottom. Scale bar 10µm. **B**) Zoom-ins of *U2OS* cells from figure 5D showing GFP-tagged Snz fragments (green) along with ER marker HSP90B1 (red) and LDs (MDH, blue). Scale bar 10µm. **C**) Schematic of CRISPR method to remove *SNZ* locus. **D**) FB tissue expressing mCD8-GFP (green) (*Dcg-Gal4>UAS-mCD8-GFP*) from control (*Dcg-Gal4* alone) or *SNZ*^RNAi^-treated FB (*Dcg-Gal4>UAS-SNZ*^*RNAi*^). Arrows indicate regions of PM “bubbling” due to PM wrapping around large pLDs. Scale bar 10µm. **E**) Total TAG in fed L3 larvae and fasted larvae for 12 hr and 24 hr periods in *wildtype* (black) and *SNZ*^KO^ larvae (blue). The lighter black-to-gray and blue-to-light blue indicates time over the fasting period. **F**) Total free fatty acids (FFAs) at fed state and during fasting for 12 hr and 24 hr periods in wildtype (black) and *SNZ*^KO^ larvae (blue).

**SFigure 6: Tissue-specific Snz over-expression promotes TAG storage in a DESAT1 dependent manner. A**) pLD size distribution for pLDs in *wildtype* control (*Dcg-Gal4*) and FB-specific transgenic Snz over-expression (Snz-Tg, *Dcg-Gal4>UAS-Sz-GFP*). **B**) mLD size distribution in wildtype control (*Dcg-Gal4*) and Snz over-expression in the FB (Snz-Tg). **C**) Thin layer chromatography (TLC) plate showing TAG levels in control (*Dcg-Gal4*) and Snz-Tg flies over time during total food deprivation assay as in Figure 6E. **D**) Survival plot of *Drosophila* flies over their lifetime fed either a chronic normal diet of 5% sugars (solid lines, see Methods) or a diet supplemented with 35% sucrose (high sugar diet, dotted lines). Control (black, *Dcg-Gal4*) or tissue-specific Snz-GFP over-expression in the FB (red, *Dcg-Gal4>UAS-Snz*, Snz-Tg) are compared. **E**) Table of major hit genes-of-interest from comparative mass spectrometry-based proteomics of Snz-GFP immuno-precipitated from Snz-Tg L3 larvae. PSM: peptide spectrum matches. **F**) Total fly free fatty acid (FFA) levels in control (*Dcg-Gal4* alone), Snz-Tg (*Dcg-Gal4>UAS-Snz*), *DESAT1*^*RNAi*^ in the FB (*Dcg-Gal4>DESAT1*^*RNAi*^), or Snz-Tg with *DESAT1*^*RNAi*^ (*Dcg-Gal4>UAS-Snz-GFP; Dcg-Gal4>UAS-DESAT1*^*RNAi*^). * indicates p<0.05. **G**) Total larval free fatty acid (FFA) levels in control (*Dcg-Gal4* alone) or conditions as in Panel F. * indicates p<0.05. **H**) Schematic of saturated (sat) fatty acid (FA) processing into unsaturated (unsat) FAs and their incorporation into TAG through the DGAT TAG synthase enzyme. Snz-GFP-interacting enzymes Bubblegum (BGM) and DESAT1 are depicted in green with Snz. **I**) Mass spectrometry-based Lipidomic Profiling of TAG-associated acyl chains by chain length and saturation number. Samples are L3 larvae from control (*Dcg-Gal4 alone*) and Snz-Tg (*Dcg-Gal4>UAS-Snz*). Samples are plotted as relative abundance of FAs. * indicates p<0.05.

**SMovie 1**: Z-section confical stack of *Drosophila* FB tissue expressing mCD8-GFP (green) and stained for LDs (MDH, magenta). In the cell periphery small pLDs are visible and are closely associated with PM wrapping. Scale bar 10um.

