## Supplementary material for "*Drosophila* Snazarus regulates a lipid droplet population at plasma membrane-droplet contacts in adipocytes": SFigures

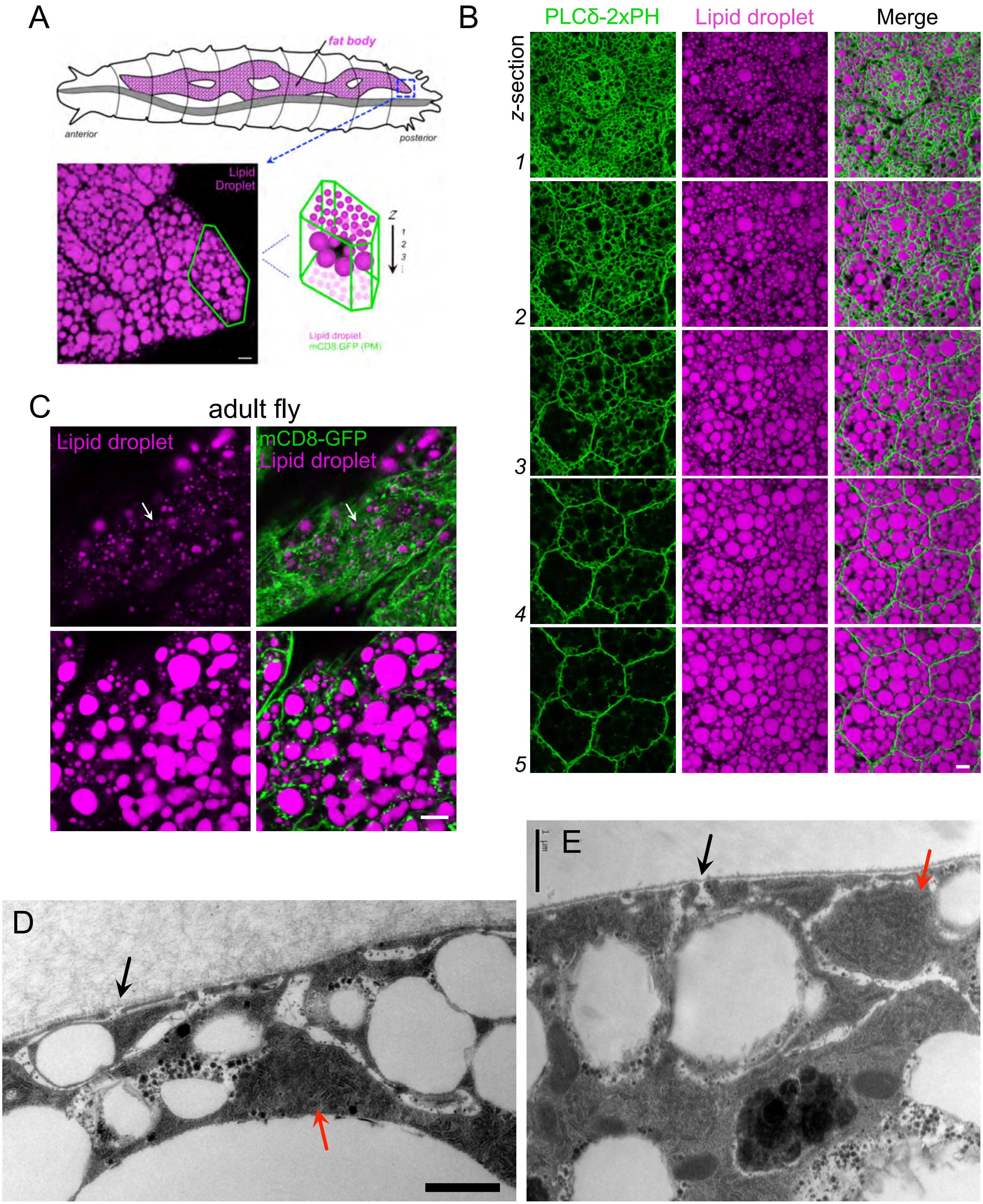
A
fat body
anterior
posterior
Lipid Droplet
Z
1
2
3
Lipid droplet
mCD8:GFP (PM)
B
PLCδ-2xPH
Lipid droplet
Merge
z-section
1
2
3
4
5
C
adult fly
Lipid droplet
mCD8-GFP
Lipid droplet
D
E

SFigure 1

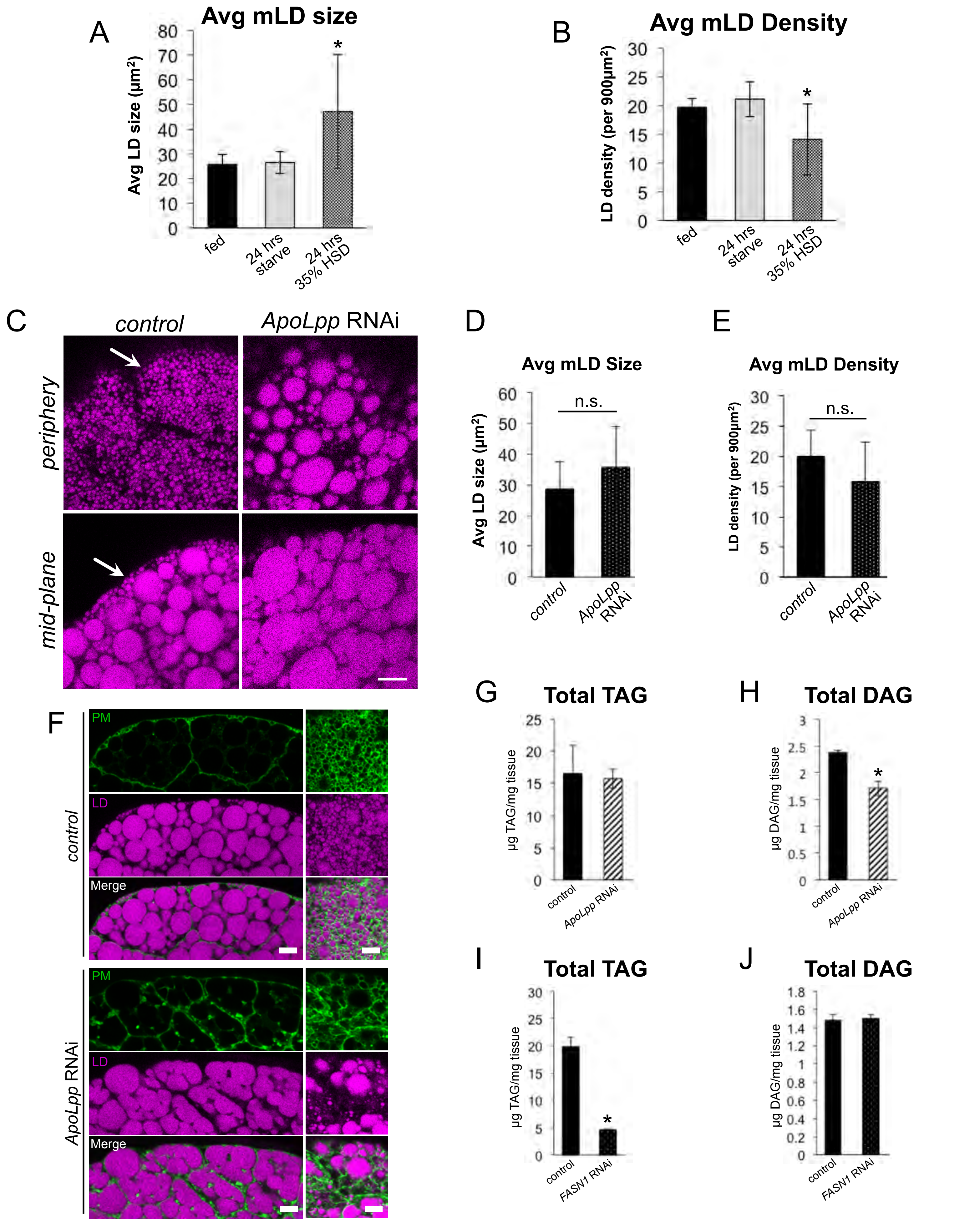

A
Avg mLD size
Avg LD size (μm²)
fed
24 hrs starve
24 hrs 35% HSD
B
Avg mLD Density
LD density (per 900μm²)
fed
24 hrs starve
24 hrs 35% HSD
C
control
ApoLpp RNAi
periphery
mid-plane
D
Avg mLD Size
Avg LD size (μm²)
n.s.
control
ApoLpp RNAi
E
Avg mLD Density
LD density (per 900μm²)
n.s.
control
ApoLpp RNAi
F
PM
LD
Merge
control
ApoLpp RNAi
G
Total TAG
μg TAG/mg tissue
control
ApoLpp RNAi
H
Total DAG
μg DAG/mg tissue
control
ApoLpp RNAi
I
Total TAG
μg TAG/mg tissue
control
FASN1 RNAi
J
Total DAG
μg DAG/mg tissue
control
FASN1 RNAi


SFigure 2

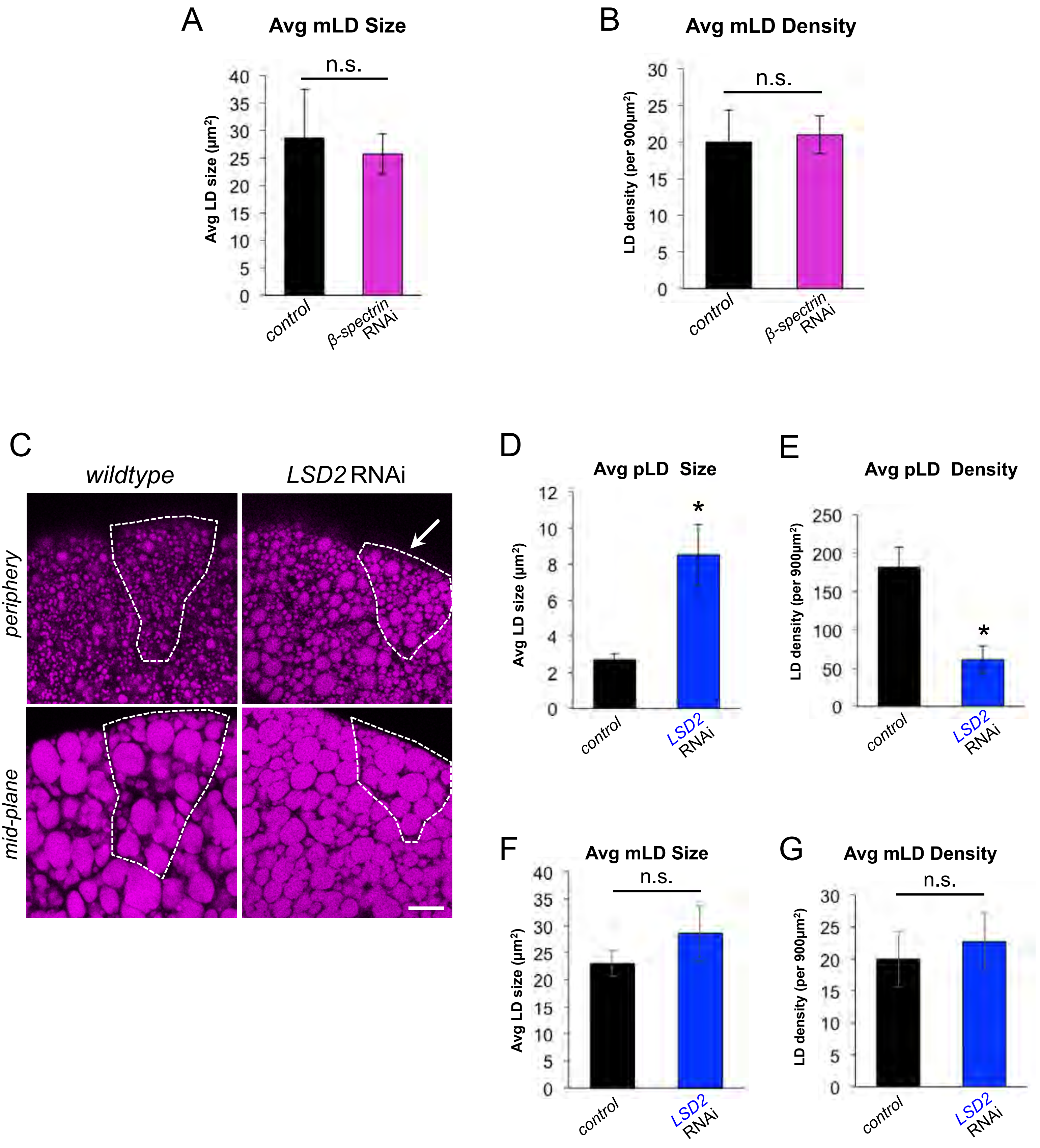
A
Avg mLD Size
n.s.
Avg LD size (µm²)
control
β-spectrin RNAi
B
Avg mLD Density
n.s.
LD density (per 900µm²)
control
β-spectrin RNAi
C
wildtype
LSD2 RNAi
periphery
mid-plane
D
Avg pLD Size
Avg LD size (µm²)
control
LSD2 RNAi
E
Avg pLD Density
LD density (per 900µm²)
control
LSD2 RNAi
F
Avg mLD Size
n.s.
Avg LD size (µm²)
control
LSD2 RNAi
G
Avg mLD Density
n.s.
LD density (per 900µm²)
control
LSD2 RNAi

SFigure 3

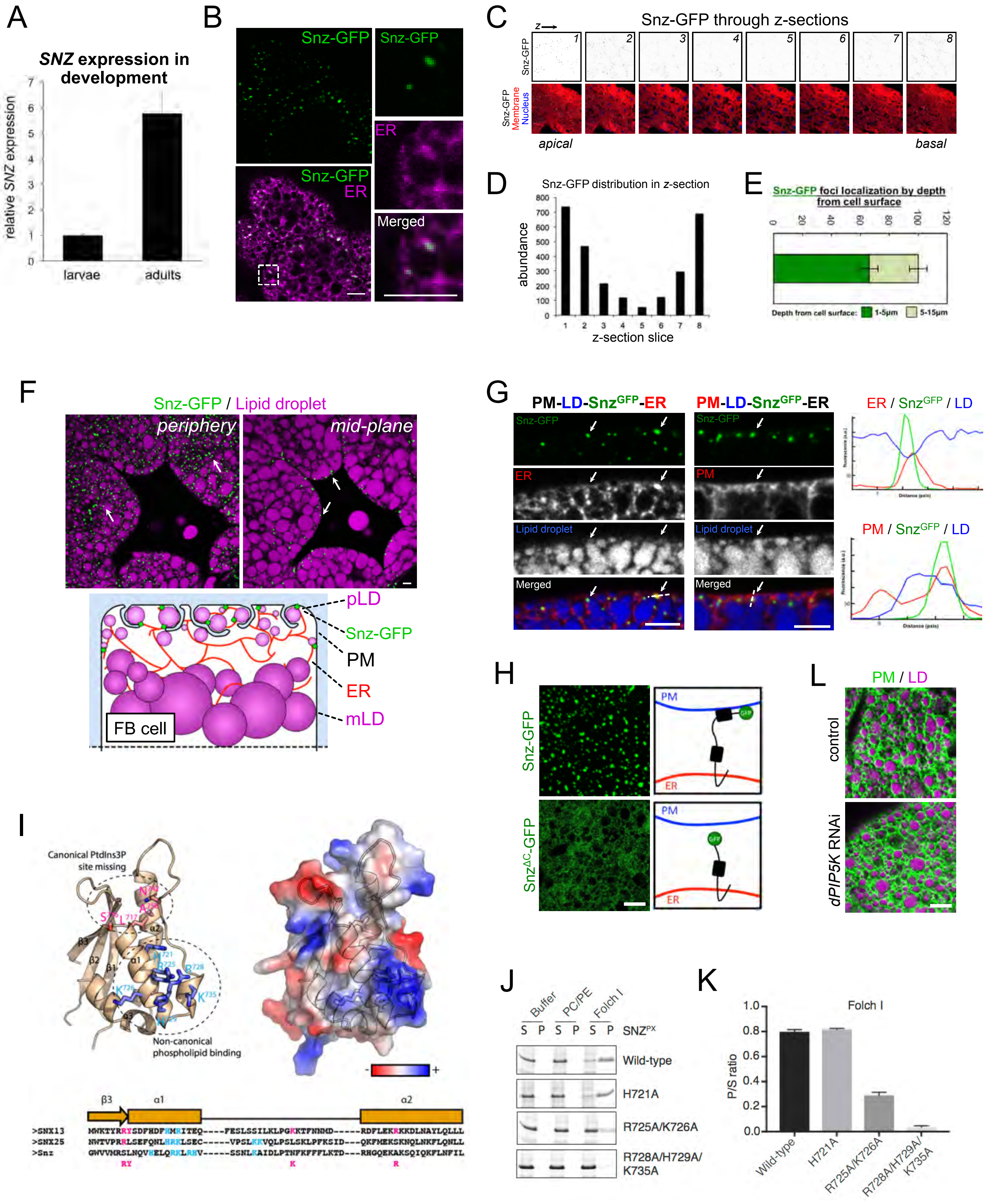
A
SNZ expression in development
relative SNZ expression
larvae
adults
B
Snz-GFP
ER
Merged
C
Snz-GFP through z-sections
apical
basal
D
Snz-GFP distribution in z-section
abundance
z-section slice
E
Snz-GFP foci localization by depth from cell surface
Depth from cell surface: 1-5μm 5-15μm
F
Snz-GFP / Lipid droplet
periphery
mid-plane
pLD
Snz-GFP
PM
ER
mLD
FB cell
G
PM-LD-SnzGFP-ER
Lipid droplet
ER / SnzGFP / LD
PM / SnzGFP / LD
H
Snz-GFP
SnzΔC-GFP
PM
ER
L
PM / LD
control
dPIP5K RNAi
I
Canonical PtdIns3P site missing
Non-canonical phospholipid binding
J
Buffer
PC/PE
Folch I
S P S P S P SNZPX
Wild-type
H721A
R725A/K726A
R728A/H729A/ K735A
K
Folch I
P/S ratio
Wild-type
H721A
R725A/K726A
R728A/H729A/ K735A

SFigure 4

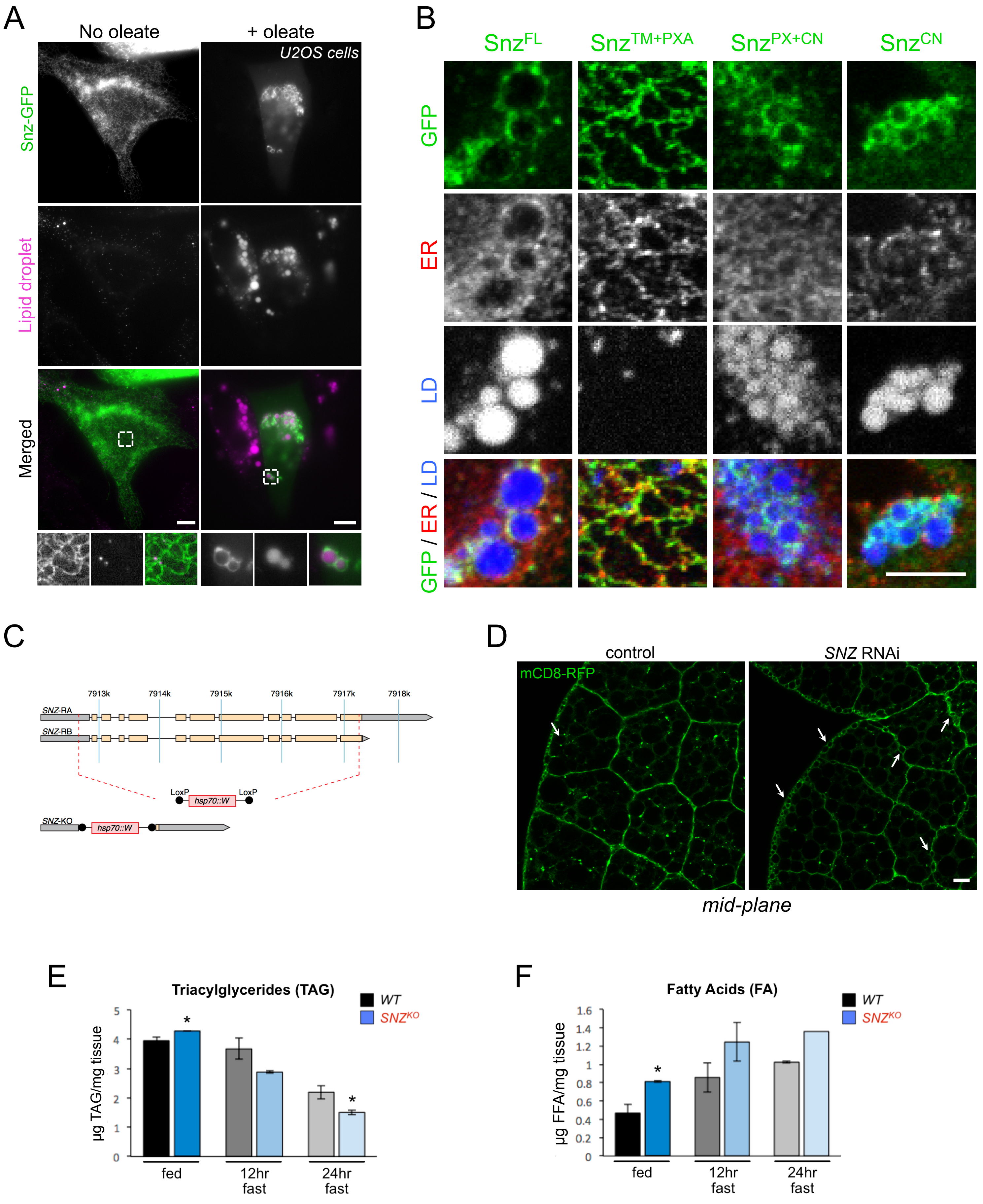
A
No oleate
+ oleate
U2OS cells
Snz-GFP
Lipid droplet
Merged
B
SnzFL
SnzTM+PXA
SnzPX+CN
SnzCN
GFP
ER
LD
GFP / ER / LD
C
7913k
7914k
7915k
7916k
7917k
7918k
SNZ-RA
SNZ-RB
LoxP
hsp70::W
LoxP
SNZ-KO
hsp70::W
D
control
SNZ RNAi
mCD8-RFP
mid-plane
E
Triacylglycerides (TAG)
WT
SNZKO
µg TAG/mg tissue
fed
12hr fast
24hr fast
F
Fatty Acids (FA)
WT
SNZKO
µg FFA/mg tissue
fed
12hr fast
24hr fast

SFigure 5

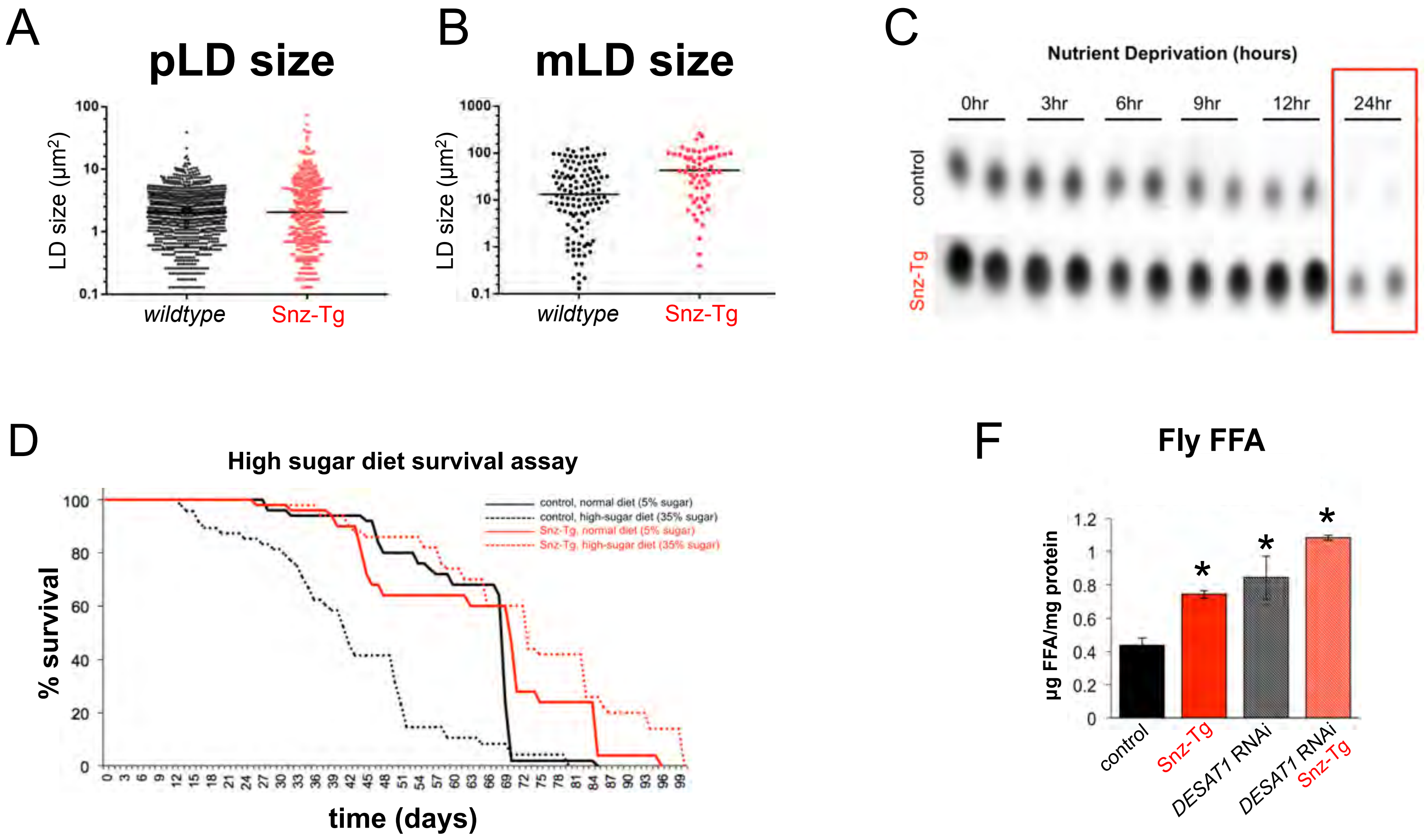


E

| Proteomics Analysis of Snz-GFP immuno-precipitation | | | | |
|---|---|---|---|---|
| Protein | PSMs | % Coverage | Spectral Index fold-enrichment (Snz-GFP/GFP) | Function |
| Desat1 | 18 | 24 | 431-fold | fatty acid desaturase; converts saturated FAs to unsaturated FAs |
| Bgm | 15 | 14.1 | 333-fold | fatty acyl-CoA ligase; generates FA-CoA from FFA |
| Cyp4e2 | 5 | 8.9 | 6.4-fold | Cytochrome P450 protein implicated in FA desaturation |
| Myosin HC | 29 | 58 | 207-fold | Actin-binding motor protein |
| Gelsolin | 14 | 7.8 | 731-fold | Actin-binding protein involved in cortical actin remodeling |

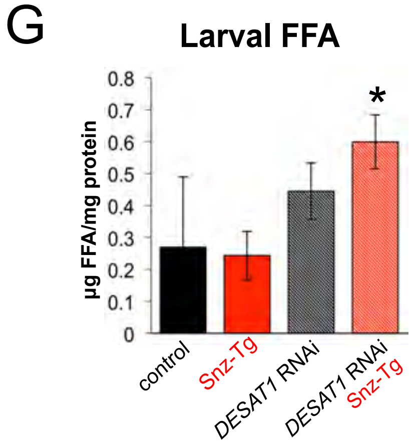


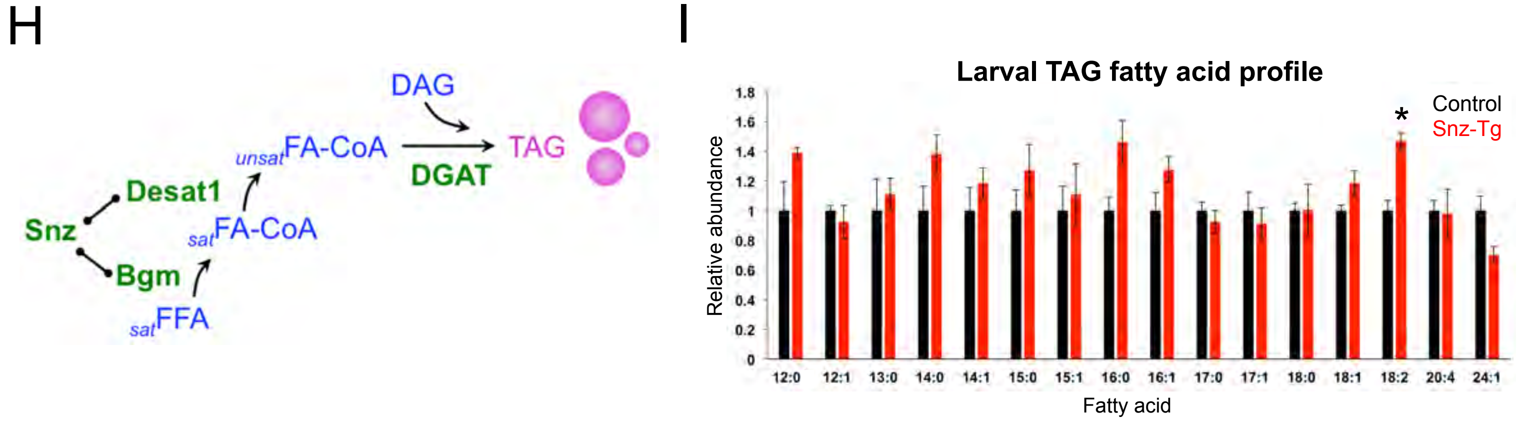


SFigure 6
